# Single-cell resolution of MET- and EMT-like programs in osteoblasts during zebrafish fin regeneration

**DOI:** 10.1101/2021.04.29.440827

**Authors:** W. Joyce Tang, Claire J. Watson, Theresa Olmstead, Christopher H. Allan, Ronald Y. Kwon

## Abstract

Zebrafish regenerate fin rays following amputation through epimorphic regeneration, a process that has been proposed to involve the epithelial-to-mesenchymal transition (EMT). We performed single-cell RNA-sequencing (scRNA-seq) to elucidate osteoblastic transcriptional programs during zebrafish caudal fin regeneration. We show that osteoprogenitors are enriched with components associated with EMT and its reverse, mesenchymal-to-epithelial transition (MET), and provide evidence that the EMT markers *cdh11* and *twist2* are co-expressed in dedifferentiating cells at the amputation stump at 1 dpa, and in differentiating osteoblastic cells in the regenerate, the latter of which are enriched in EMT signatures. We also show that *esrp1*, a regulator of alternative splicing in epithelial cells that is associated with MET, is expressed in a subset of osteoprogenitors during outgrowth. This study provides a single cell resource for the study of osteoblastic cells during zebrafish fin regeneration, and supports the contribution of MET- and EMT-associated components to this process.

## INTRODUCTION

While humans have limited potential for limb regeneration, some vertebrates can regenerate bony appendages following amputation. Zebrafish can regenerate amputated caudal fins through the formation of a pool of lineage-restricted progenitors called the blastema [1]. Following fin amputation, mature osteoblasts at the amputation stump dedifferentiate, migrate to the blastema, self-renew, then redifferentiate to form new bone [2–5]. This process begins within the first 24 hours after amputation, with osteoblast redifferentiation beginning by 2 days post amputation (dpa) as detected by the upregulation of the committed osteoblast marker, *sp7*, followed by the expression of the late differentiation marker *bglap* detectable by 5-6 dpa. In addition to dedifferentiation of mature cells, osteoprogenitors also derive from resident mesenchymal precursors within the fin ray joint [6]. The formation of osteoprogenitors through dedifferentiation in zebrafish contrasts from mammals, where osteoblasts primarily derive from resident precursors. The cell biological processes that enable dedifferentiation and reversible switching between differentiated and self-renewing cell states are poorly understood.

Cell plasticity and reversibility between metastable cell states are hallmarks of the epithelial-to-mesenchymal transition (EMT), and its reverse, mesenchymal-to-epithelial transition (MET). EMT is a developmental program by which cells lose epithelial (E) characteristics (e.g., E-cadherin, desmosomes, cortical actin, apical–basal polarity) while adopting features of mesenchymal (M) cells (e.g., increased vimentin) [7]. EMT is often observed during cancer invasion and metastasis, where it results in dedifferentiation and acquisition of cancer stem cell-like properties [8]. While cancer progression is impaired by complete EMT, cancer cell stemness is enhanced by partial EMT, where cells reversibly switch between metastable cell states with hybrid E/M phenotypes that are epithelial-shifted or mesenchymal-shifted [9]. Epimorphic regeneration exhibits a number of biological parallels to the progression of solid tumors. It has been suggested that certain mechanisms enabling regeneration may be co-opted by cancer to promote growth at primary and metastatic sites [10]. *In silico* evidence of a sequential EMT and MET process occurring during axolotl limb regeneration has been reported [11], however we are unaware of other articles reporting the existence of EMT signatures in epimorphic regenerative models. A review by Wong and Whited specifically states that no EMT signatures were identified in two axolotl limb regeneration scRNA-seq studies, indicating that more investigation is needed [10, 12, 13].

There is evidence that mammalian osteoblastic cells differentially exhibit characteristics of mesenchymal and epithelial cells, which may be conserved in zebrafish. In mammals, early osteoblastic cells are stromal cell-like in appearance, whereas matrix-secreting osteoblasts are tall or cuboidal and form a continuous layer, bearing a partial resemblance to epithelial cells [14]. The primary form of cell-cell adhesion between osteoblasts is through adherens junctions [15, 16], which are typically found in epithelial and endothelial cells and rely on the interaction of cadherins at the cell surface to form an intercellular adhesion complex. Multiple studies have shown that interfering with cadherin function impairs osteoblast differentiation [17]. Upon further maturation, active osteoblasts transition into quiescent bone lining cells, appearing as flat and elongated cells on the mineralized surface. Thus, while mesenchymal in origin, mammalian osteoblastic cells differentially exhibit characteristics of mesenchymal and epithelial cells depending on cell state.

During zebrafish fin regeneration, Stewart et al., 2014, noted dual mesenchymal and epithelial characteristics in osteoblastic cells, and hypothesized this was due to EMT [18]. Specifically, they observed that blastemal osteoprogenitors expressed the EMT-associated transcription factors (EMT-TFs) *twist2* and *twist3*, loss of the epithelial marker a-catenin 1, and changed from an elongated to a polygonal morphology. However, whether this reflects an EMT-like transition or a function independent of EMT is poorly understood. Only a limited set of EMT markers have been characterized, which may be insufficient to resolve hybrid cells that exhibit both E and M characteristics [7]. Moreover, multiple markers are often needed to precisely assess the state of a cell undergoing EMT within specific biological contexts [7]. Finally, many transcription factors and other components critical for EMT also affect other cellular functions. These components may be regulated through programs that involve canonical EMT [7]. As such, more in-depth molecular characterizations are needed. While single cell analyses are comprehensive approaches and can provide valuable insight, previous single-cell RNA-seq (scRNA-seq) studies of zebrafish caudal fin regeneration have reported low numbers of osteoblastic cells [19], hampering an in-depth analysis.

Here, we performed sci-RNA-seq (single-cell combinatorial indexing RNA sequencing) [20] to elucidate osteoblastic transcriptional programs during zebrafish caudal fin regeneration. We speculated that the higher throughput of sci-RNA-seq in concert with differences in sample preparation could help retrieve osteoblastic cells compared to droplet-based scRNA-seq approaches. Specifically, sci-RNA-seq uses combinatorial indexing for transcriptomic profiling, enabling higher throughput compared to droplet-based scRNA-seq where individual cells are isolated within physical compartments [20]. Moreover, whereas droplet-based scRNA-seq commonly employs tissue dissociation using enzymes and usually heat [21], sci-RNA-seq commonly employs manual homogenization, which could help retrieve cells easily destroyed by or not readily liberated by enzymatic tissue dissociation. Thus, in contrast to existing scRNA-seq resources [19], our dataset captures a high number of osteoblastic cells (1,673 nuclei), allowing for their in-depth analysis. We show that osteoprogenitors are enriched with components associated with the epithelial-to-mesenchymal transition (EMT) and its reverse, mesenchymal-to-epithelial transition (MET). In trajectory analyses, osteoblastic cells solely expressed EMT components, or transiently expressed MET components prior to expressing those for EMT. We provide evidence that the EMT markers *cdh11* and *twist2* [22] are co-expressed in dedifferentiating cells at the amputation stump at 1 dpa and in differentiating osteoblastic cells in the regenerate at 3 dpa, and that EMT signatures are enriched in the latter. We also show that *esrp1*, a regulator of alternative splicing in epithelial cells whose expression is important for MET [23], is expressed in a subset of osteoprogenitors during outgrowth. This study provides a single cell resource for the study of osteoblastic cells during zebrafish fin regeneration, and supports the contribution of MET- and EMT-associated components to this process.

## RESULTS

### sci-RNA-seq captures rare osteoblastic populations

We performed sci-RNA-seq using 225 fins from 3-5 days post amputation (dpa), which yielded 19,603 nuclei after filtering for quality control. At these timepoints, a variety of tissues have regenerated (Fig 1A). Unbiased clustering identified 12 different cell populations, of which we defined 10 out of 12 by cell/tissue type using published biomarkers (Fig 1BC, Table S1) [2, 5, 19, 24–47]. All populations were observed at both 3 and 5 dpa (see Figure S1A). There was noticeable variability in the proportion of cell types in two different samples from 3 dpa (see Figure S1BC), indicating that the relationship between regeneration timepoint and cell type proportion could not be reliably inferred. Thus, all timepoints were pooled for final analysis.

**Figure 1.**
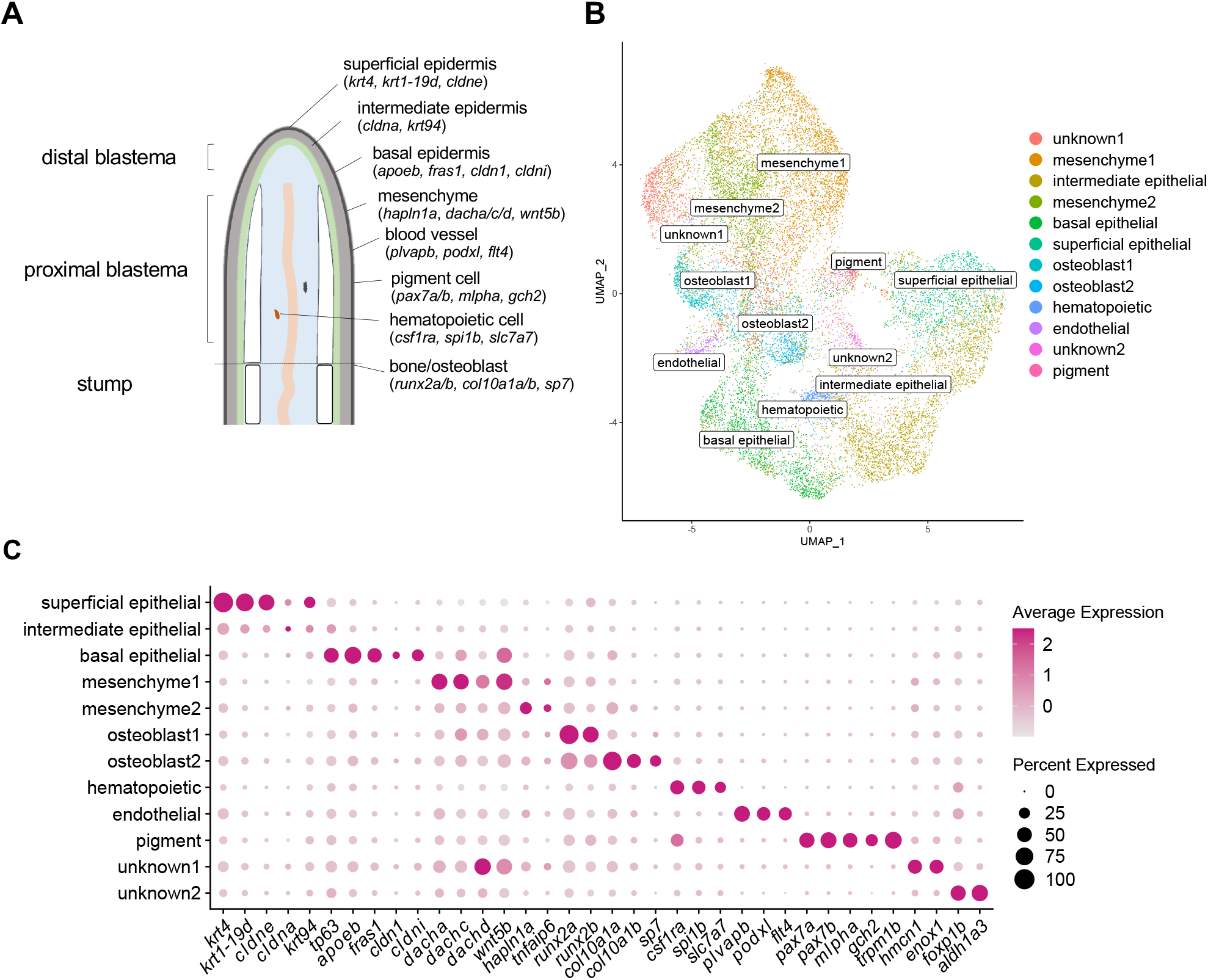
Molecular profiling of fin regeneration reveals distinct cell types including two osteoblast populations. (A) Schematic of zebrafish fin regeneration showing tissues and corresponding gene markers. (B) UMAP plot visualizing 19,603 cells from sci-RNA-seq analysis of regenerated fins at 3 and 5 dpa. Of the 12 distinct clusters, 10 tissue types could be identified by molecular markers, including two osteoblast populations. (C) Dotplot visualizing expression of genes demarcating the 12 cell clusters. Circle size represents the percentage of cells expressing the gene, and color indicates average gene expression. See also Figure S1.

We identified two populations of osteoblastic cells—osteoblast1 (973 cells) and osteoblast2 (700 cells)— marked by the strong expression of the osteoprogenitor marker *runx2a/b* (Fig 1C), a transcription factor expressed during early osteoblast differentiation. Mapping of osteoblast1 and osteoblast2 identities to a previous scRNA-seq dataset of fin regeneration [19] revealed 16 and 23 osteoblast1 and osteoblast2 cells, respectively (~40x fewer cells), indicating the difficulty of capturing osteoblastic cells in scRNA-seq studies.

Osteoprogenitors can differentiate into osteoblasts and joint-forming cells [48] (Fig 2A). Cells within the osteoblast1 cluster expressed genes known to be upregulated during osteoprogenitor differentiation into joint cells. This included *pthlha*, a marker of joint cells [7], and *mmp9*, a marker of joint-derived resident osteoprogenitors [6] (Fig 2B). In contrast, the osteoblast2 cluster expressed genes known to be specifically expressed during osteoblast differentiation, notably *col10a1a/b* (a marker of pre-osteoblasts), *sp7* (a marker of immature osteoblasts), and *ihha* [7] (Fig 2B). The osteoblast1 and osteoblast2 clusters scored higher for joint cell and osteoblast gene signatures, respectively (Fig 2B). We identified genes with significant differences in expression between the two osteoblast groups (see Table S2). These genes included *lmx1ba* in the osteoblast1 cluster (Fig 2C), and *slc8a4b* in the osteoblast2 cluster (Fig 2D). We also identified genes with significantly increased expression in both groups, compared to non-osteoblastic cell types (see Table S3). This included *cdh11* (Fig 2E), which was recently identified by scRNA-seq to be a potential marker of osteoblastic cells, but still remains to be experimentally validated [19].

**Figure 2.**
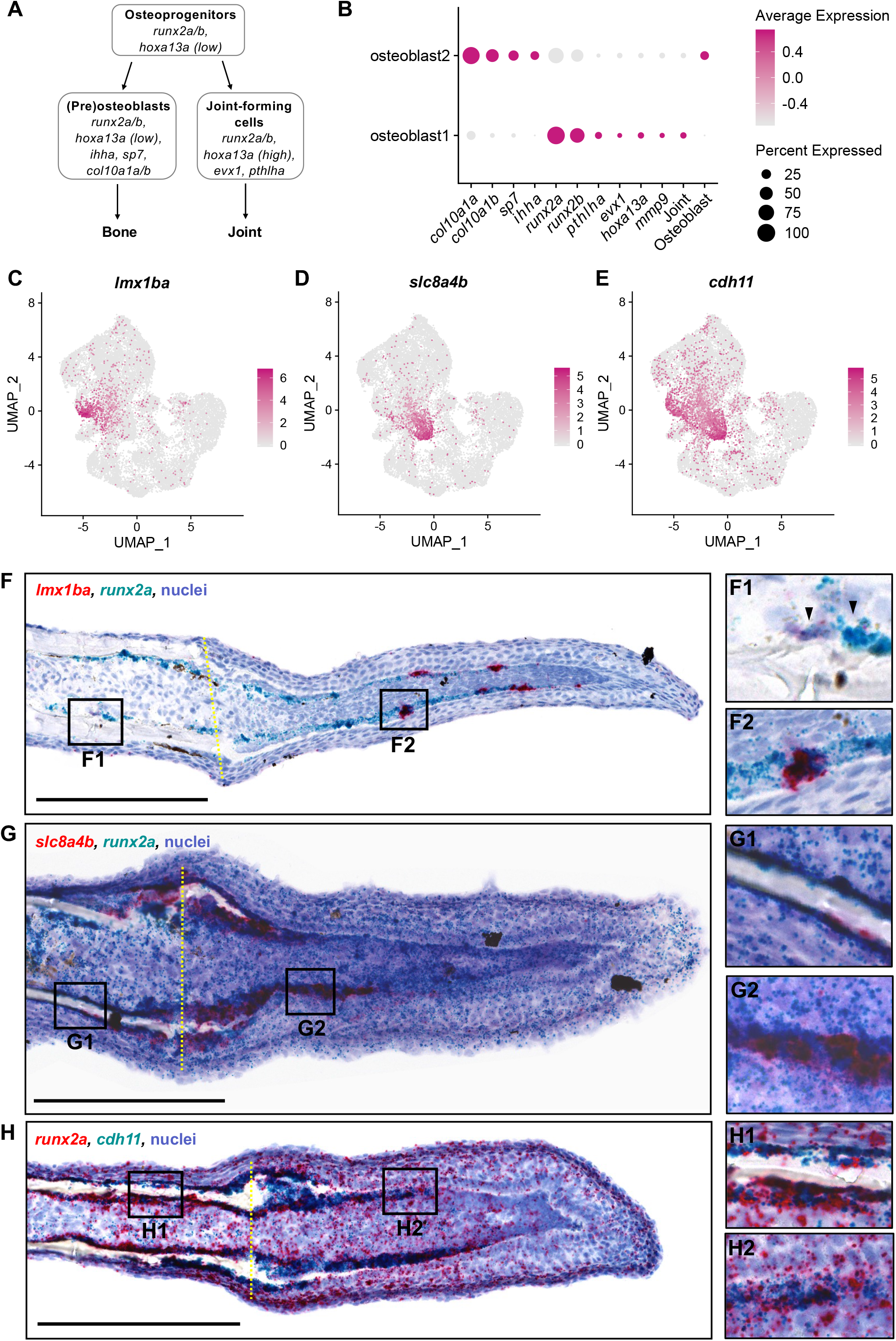
Osteoblast1 and osteoblast2 are likely enriched with cells fated to become joint cells and osteoblasts, respectively. (A) Schematic of osteoprogenitor differentiation into either osteoblasts or joint-forming cells and corresponding gene markers (adapted from McMillan et al., 2018). (B) Dotplot comparing the expression of joint cell markers and bone markers in the two osteoblast populations. The clusters osteoblast1 and osteoblast2 are enriched with genes associated with joint-forming cells and osteoblasts, respectively. (C-E) Feature plots for *lmx1ba*, *slc8a4b*, and *cdh11*. (F-H) Dual RNA in situ hybridization in 3 dpa fin cryosections for the indicated genes. All scale bars are 200 μm. Amputation planes are indicated with a yellow dotted line. (F) *lmx1ba*+/*runx2a*+ cells were present in cells near the native joint (F1 inset, black arrowheads), and in presumptive joint cells in the regenerate (F2 inset). (G) *slc8a4b*+/*runx2a*+ cells were present at the amputation stump, near the regenerating bone, and in cells along the proximal lateral mesenchymal compartment. (H) *cdh11*+ cells lined the native bone (H1 inset), the regenerating bone near the amputation stump, and cells along the lateral edges of the mesenchymal compartment of the blastema (H2 inset).

Expression of *lmx1ba, slc8a4b*, and *cdh11* has not previously been characterized during zebrafish fin regeneration. Thus, we performed dual RNA *in situ* hybridization (ISH) to stain for transcripts for these genes in concert with *runx2a*. Previous fin regeneration studies have shown that at 3 dpa, osteoprogenitors in the fin blastema are located distal to the amputation stump, extending along the lateral edges of the mesenchymal compartment, just deep to the basal epidermis. We found high expression of *runx2a* in cells residing in these areas. Although we did not do lineage analysis in our studies, based on their location within the blastema, cells in these areas that were strongly expressing *runx2a* were most likely to be osteoprogenitors. We also observed lower expression of *runx2a* in the epidermis and mesenchyme. Our scRNA-seq data indicate that *runx2a* is expressed in some cells within the epidermis and mesenchyme populations (see Figure 1C). While this was not the focus of this study, our data support the notion of low expression of *runx2a* in epidermis and mesenchyme during fin rengeneration.

We observed staining for the osteoblast1 marker *lmx1ba* in distinct, relatively evenly spaced clusters of *runx2a+* cells along the lateral mesenchymal compartment of the blastema (Fig 2F2), similar to the staining pattern of the known joint markers *pthlha, evx1*, and *hoxa13a* [48]. We also detected *runx2a*+/*lmx1ba*+ cells in the native fin ray joints (Fig 2F1, black arrowheads). These data support the potential for *lmx1ba*+ cells in the blastema to be presumptive regenerating joint cells of the fin rays. Mutations in the human ortholog *LMX1B* have been implicated in focal segmental glomerulosclerosis (a disease of the kidney) and nail-patella syndrome, which is commonly manifested by underdeveloped or irregularly shaped nails, elbows, and knees [49]. *Lmx1b*-deficient mice exhibit abnormalities in dorsal limb structures attributable to defects in dorsal-ventral patterning [50]. In mice, *Lmx1b* appears to be expressed between, rather than within, developing bones [51] and it has been postulated that *Lmx1b* could have additional function in anterior-posterior patterning [49]. Further studies of the function of *lmx1ba* in zebrafish fin regeneration is warranted.

Next, we examined the osteoblast2 marker *slc8a4b* (Fig 2G). We observed staining for *slc8a4b* in *runx2a+* cells lining the distal end of the amputation stump, around the newly regenerating bone, and in cells along the proximal lateral mesenchymal compartment where new bone was likely to form (Fig 2G2). Staining for *slc8a4b* was largely absent in *runx2a+* cells in the native bone (Fig 2G1). These data support the potential for *slc8a4b+* cells to be a population of osteoblast progenitors. *slc8a4b* was first identified during a screen for homologs of the mammalian genes *Slc8a1/2/3* (formerly *NCX1/2/3)*, which are members of the calcium/cation antiporter family of proteins that help regulate intracellular calcium in a variety of cell types. Initially called NCX4, *slc8a4b* was shown to be a distinct fourth isoform of the NCX family, and orthologs of *NCX4/slc8a4b* have only been identified in bony fishes [52]. In our studies, a tBLASTn of *slc8a4b* showed highest similarity to mammalian *Slc8a1* (XM_006523943.2, E-value: 0, 96% query cover, 68% identity). In humans, genetic variants near *SLC8A1* have been associated with risk of bone fracture [53]. In mice, *Slc8a1* is expressed in osteoclasts and plays a role in bone resorption [54], however its function in bone formation requires further study.

Finally, we examined expression of *cdh11* (Fig 2H). We observed prominent staining for *cdh11* in *runx2a*+ cells lining the amputation stump, in the lateral mesenchymal compartment (Fig 2H2), and in the native bone (Fig 2H1). *CDH11* is highly expressed in osteoblastic cell lines and upregulated during differentiation. *Cdh11*-deficient mice exhibit hypoplastic synovium [55], resistance to inflammatory arthritis [55], and osteopenia [56]. These studies support the notion that osteoblast1 and osteoblast2 cells are distinct populations enriched for cells fated to become joint cells and osteoblasts, respectively.

### Osteoblastic cells exhibit distinct cell states and trajectories

We subset the osteoblast2 cells for further analysis. Unbiased clustering of the osteoblast2 cells revealed five subclusters (Fig 3A). Analysis of markers of osteoblast differentiation (Fig 3B) and cell cycle phase (Fig 3C) revealed distinct cell cycle and differentiation states. Clusters 4 and 5 had the highest osteoblast differentiation scores. Whereas cluster 5 was enriched for *sp7*, cluster 4 was enriched for *col10a1a/b*. Moreover, cluster 5 had the lowest percentage of cells in the G2/M and S phases. Cluster 3 had a lower osteoblast differentiation score compared to clusters 4 and 5. Moreover, these cells had relatively high expression of *col10a1a/b*, and had the highest percentage of cycling cells as indicated by cells in the G2/M and S phases. Cluster 3 also had low *runx2a/b* expression, which is in agreement with previous studies showing that murine RUNX2 levels regulate cell cycle entry and exit in osteoblasts, with minimal levels of RUNX2 during G2 and mitosis [57, 58]. Clusters 1 and 2 had the lowest osteoblast differentiation scores, and had higher *runx2a/b* expression compared to clusters 3 and 4. In regard to statistical analysis, using the Wilcoxon rank sum test with Bonferroni correction (see Figure S2) we found that for cluster 5, the osteoblast differentiation score was significantly different from clusters 1 and 2; for cluster 4, the osteoblast differentiation score was significantly different from clusters 1, 2, and 3. For cell cycle scores, using a difference of proportions test (prop.test in R) with Bonferroni correction (see Figure S3), we found that the proportion of cycling cells was significantly different between most cluster pairwise comparisons except between clusters 1 and 2, clusters 1 and 4, and clusters 2 and 4. To assess whether specific clusters were enriched with doublets, we used the computeDoubletDensity function from the scDblFinder package in R [59]. This function generates simulated doublets by randomly pairing cells from the dataset, then calculates a doublet score for each cell based on its similarity to simulated doublets (see Figure S4). We did not find clear evidence that doublets were enriched in any individual cluster, suggesting that each cluster represented a *bona fide* subpopulation. The above studies identify diverse osteoblastic subpopulations which differ in cell cycle and differentiation status, three of which (clusters 3, 4, and 5) we could identify based on cell cycle status or known osteoblastic biomarkers.

**Figure 3.**
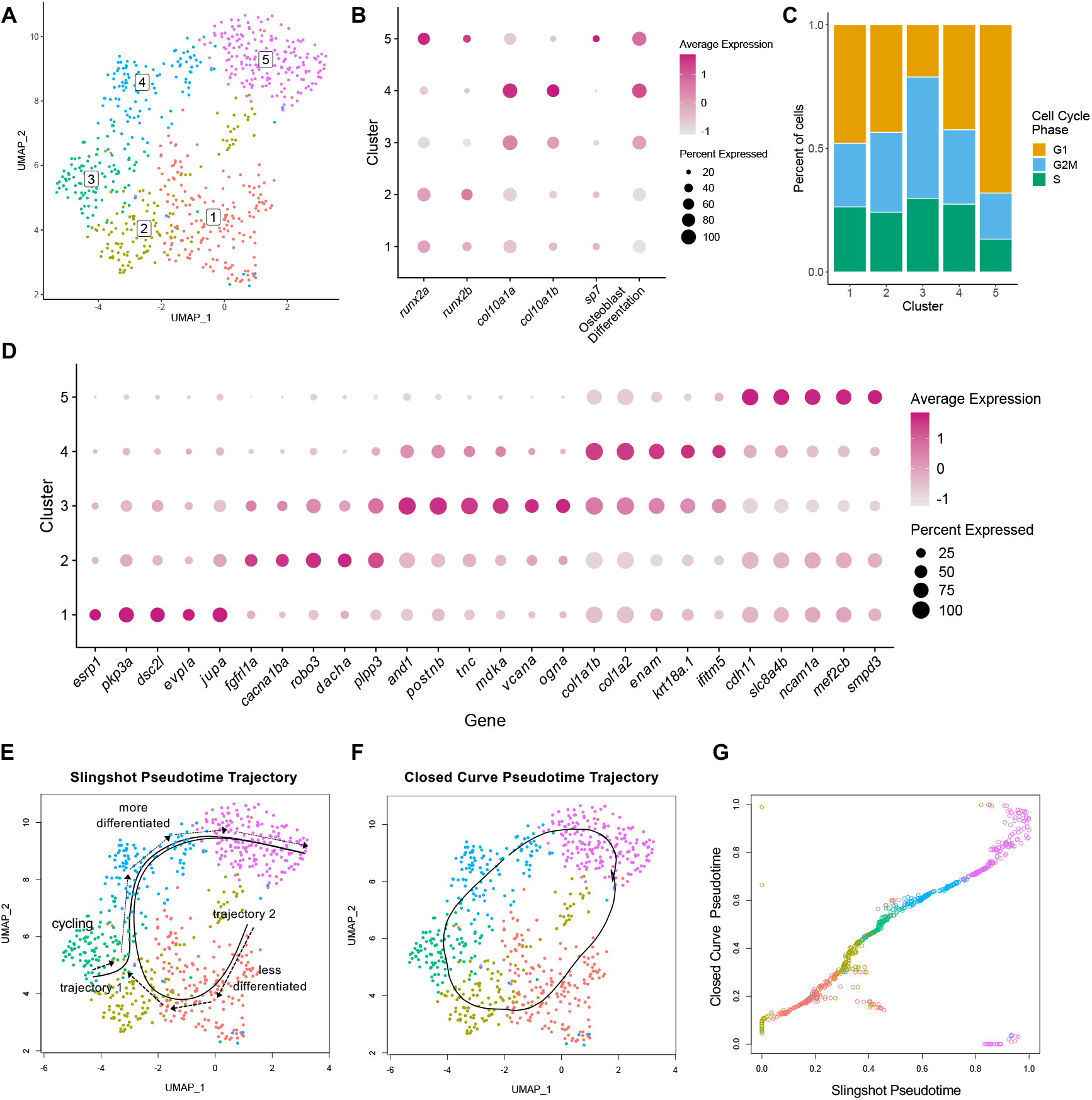
Clustering and trajectory analysis of osteoblast2 population. (A) UMAP plot showing five distinct osteoblastic subclusters. (B) Dotplot visualizing expression of known genes marking the different stages of osteoblast differentiation and osteoblast differentiation scores. (C) Percentage of cells in each cell cycle phase for each osteoblast subcluster. (D) Dotplot showing selected genes found to be differentially expressed within each osteoblastic subcluster. (E-G) Pseudotemporal ordering of osteoblast2 cells. (E) Slingshot pseudotime ordering yields two trajectories. Trajectory 1 starts in cluster 3, whereas trajectory 2 starts in cluster 1. (F) Closed curve analysis suggests a similar trajectory to trajectory 2 derived from Slingshot analysis. (G) Comparison of the closed curve trajectory to Trajectory 2 derived from Slingshot shows that the orderings from open (Slingshot) and closed curve analyses are highly similar.

Next, we performed differential gene expression analysis (Fig 3D, Table S4) and found that osteoblast2 subclusters were uniquely characterized by genes associated with the epithelium or involved in extracellular matrix formation or osteoblast differentiation. Of the 10 most differentially expressed genes in cluster 1, nine were associated with epithelial cells, including genes encoding for components of desmosomes (*dsc2l* and *evpla*) and adherens junctions (*pkp3a*). Among the top marker genes in clusters 3-4 were those encoding for extracellular matrix components. For instance, 10 of the top 20 genes of cluster 3 included markers such as *postnb, and1, tnc, vcana*, and *ogna*, while the top two most differentially expressed genes in cluster 4 encoded for Type I collagen (*col1a1b* and *col1a2*). Cluster 4 was also marked by the expression of *ifitm5* which has been implicated in skeletal disease [60]. The top four markers of cluster 5 included genes whose orthologs are expressed during osteoblast differentiation (*cdh11* [15], *mef2c* [61], *ncam1a* [62]) and/or are implicated in skeletal disease (*smpd3* [63]). A complete analysis of enrichment for pathways and GO terms can be seen in Figure S5.

To assess ordering through cell states, we performed pseudotime and trajectory analysis. In constructing trajectories, the direction of the trajectory may go from more to less differentiated (dedifferentiation), or less to more differentiated (differentiation). At 3-5 dpa, osteoblastic cells are predominantly undergoing differentiation rather than dedifferentiation [2]. Thus, we presumed the direction of the trajectory was from less to more differentiated. Because cluster 5 had the highest gene expression for the osteoblast marker *sp7*, we further presumed that cluster 5 was likely to be the terminal state. Ando et al., 2017, previously showed that *mmp9* is a marker of joint osteoprogenitors that contribute to fin regeneration [6]. In separate studies in which we subset all cells (including osteoblast1 and osteoblast2) that expressed either *sp7* or *runx2a*, we did not detect a clear *mmp9*-enriched cluster which would be needed to construct a trajectory from joint osteoprogenitor cells to differentiated osteoblasts. Thus, joint cells were not included in our trajectory analyses. Unbiased analysis using Slingshot [64] revealed two trajectories (Fig 3E). Trajectory 1 started in cluster 3 and progressed through clusters 4 and 5. In this trajectory, cycling osteoprogenitors transition into pre-osteoblasts that express *col10a1a/b*, and then further transition into immature osteoblasts that express *sp7*. Trajectory 2 started in cluster 1, and then progressed through a trajectory similar to trajectory 1. In both trajectories, extracellular matrix genes such as *postnb, and1, tnc, vcanb*, and *ogna* are expressed prior to or concomitantly with genes encoding for Type I collagen (*col1a1b* and *col1a2*). This is consistent with the synthesis of a transitional matrix rich in tenascin C prior to the synthesis of ECM proteins characteristic of differentiated skeletal tissues like collagen type I [65] during zebrafish fin regeneration and salamander limb regeneration. These studies point to multiple candidate trajectories underlying the identified cell states.

We then evaluated whether the trajectories exhibited characteristics of de- and redifferentiation. We hypothesized that de- and redifferentiation involves a cyclical progression through cell states which would manifest as a closed curve through gene expression space. We therefore sought to examine whether trajectory 2 represented a path through cellular de- and redifferentiation by assessing if the pseudotime orderings in the trajectory were similar to those derived from a closed curve. Algorithms for the analysis of cyclic trajectories are sparse compared to non-cyclic methods [66]. Moreover, ANGLE, the highest performing cyclic algorithm benchmarked in [66], requires 2D data embeddings. We therefore devised an algorithm for pseudotime inference that enables closed curve trajectory analysis using data embeddings of arbitrary dimension (Fig 3F). Pseudotime orderings derived from the closed curve trajectory were highly similar to trajectory 2 (Fig 3G). In addition, orderings derived using either UMAP or PCA embeddings were comparable to each other (see Figure S6). Thus, pseudotime orderings were similar whether using an open (Slingshot) or closed curve, the latter of which is predicted with a cyclical progression through cellular de- and redifferentiation states.

### Osteoblastic cells differentially express MET- and EMT-associated genes

Next, we assessed the mesenchymal and epithelial characteristics of osteoblast2 subclusters. For this, we computed mesenchymal and epithelial differentiation scores (Fig 4A). Whereas mesenchymal differentiation scores were relatively uniform across cell clusters, epithelial differentiation scores were higher in clusters 1-3 compared to clusters 4-5. As a consequence, mesenchymal scores were higher than epithelial scores in clusters 4-5. Hence, osteoblastic cells express genes associated with both mesenchymal and epithelial differentiation, which are differentially expressed depending on cell state. Because expression of epithelial and mesenchymal markers is expected in cells undergoing MET and EMT respectively, we next investigated whether osteoblast2 subclusters expressed MET and EMT components.

**Figure 4.**
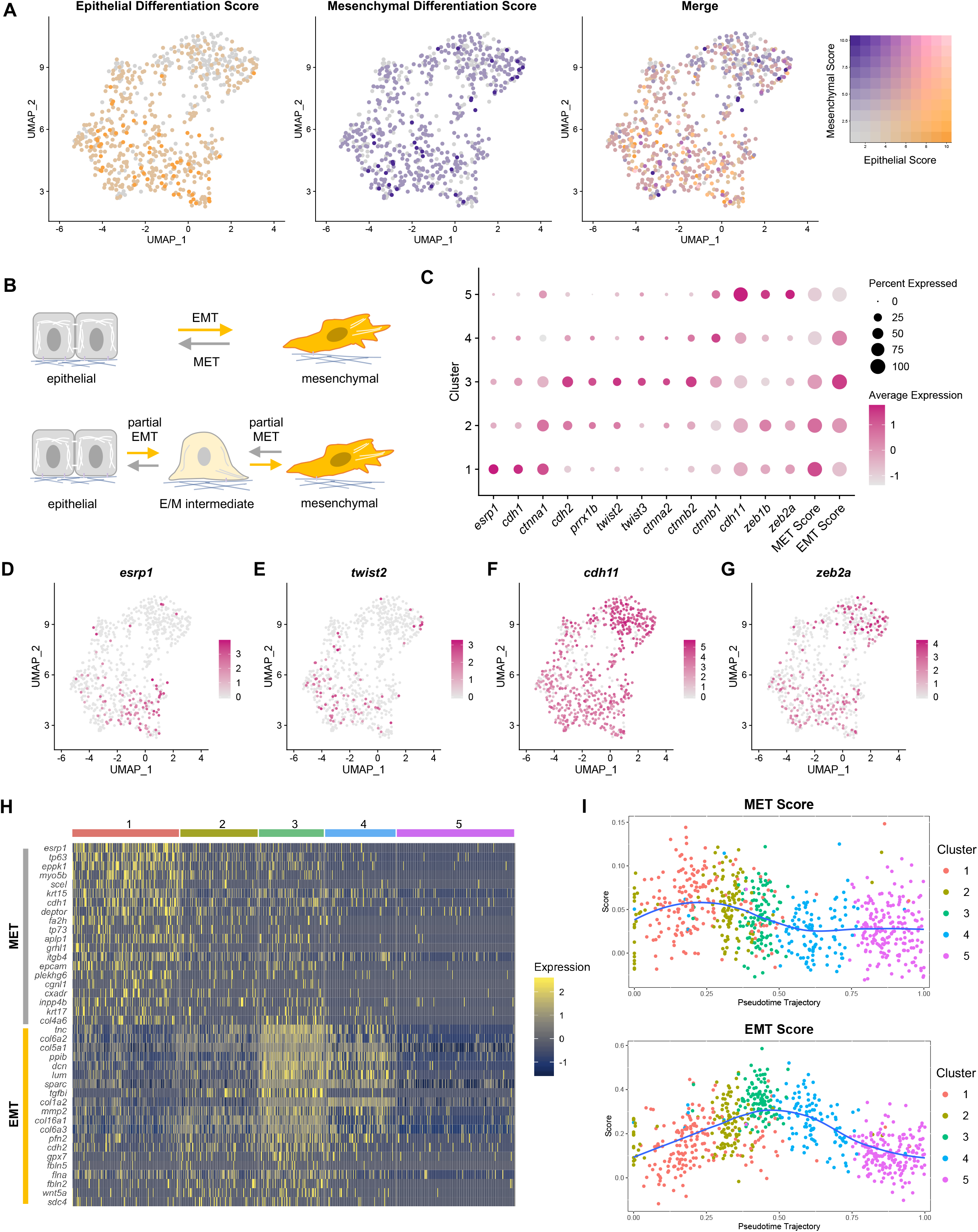
Osteoblastic cells express EMT and MET-associated genes. (A) Osteoblastic cells colored by epithelial differentiation score (left), mesenchymal differentiation score (second from left), and merged epithelial and mesenchymal differentiation scores (third from left). Color legend at right. (B) Partial EMT and MET result in intermediate cellular phenotypes that encompass dual epithelial and mesenchymal phenotypes (bottom). (C) Dotplot of the expression of canonical EMT and MET markers in the osteoblastic subclusters. Clusters were also scored for the expression of EMT and MET gene signatures. (D-G) Feature plots for *esrp1*, *twist2*, *cdh11*, and *zeb2a*. (H) Top differentially expressed MET and EMT genes in the osteoblasts as visualized by heatmap. (I) Pseudotemporal expression of MET (top) and EMT (bottom) gene signatures in the osteoblast2 cells using trajectory 2.

We first assessed whether the top markers for osteoblastic cell clusters included canonical markers for MET and EMT [22, 67–84]. We found that the second most differentially expressed gene in cluster 1 was *esrp1*. Epithelial splicing regulatory proteins (ESRP) are highly conserved RNA-binding proteins that regulate alternative splicing, and which participate in MET and EMT [85, 86]. There are two paralogs of *esrp* in vertebrates, *ESRP1* and *ESRP2*. They are required for an epithelial splicing program involving alternative splicing of hundreds of genes, many of which have functions in cell-cell adhesion, cell polarity, and cell migration [68]. This includes *Fgfr2*; *Esrp1* and *Esrp2* are required for the epithelial isoform of *Fgfr2*, and knockdown of *Esrp* results in the complete switch to the mesenchymal isoform of *Fgfr2* [85]. In zebrafish, double knockout of *esrp1* and *esrp2* leads to multiple developmental abnormalities, including malformed fins [69].

The most differentially expressed gene in cluster 5 was *cdh11* (Table S4). CDH11 is expressed during cadherin switching in EMT [87]. In this process, expression of CDH1 (epithelial cadherin) decreases in concert with increased expression of CDH11 and CDH2 (mesenchymal cadherins). As such, CDH11 serves as a biomarker of EMT [88]. In regard to its embryonic functions, CDH11 is predominantly expressed within tissues of mesodermal origin [89], and is linked to morphogenesis in the head, somite, and limb bud of early mouse embryos [80]. In addition to its expression within mesenchymal cells, *Cdh11* is also expressed in parts of the neural tube and otic vesicle in early mouse embryos [80]. In zebrafish, knockdown of Cdh11 results in otic vesicle [33] and retinal defects [90]. In chick, CDH11 expression is observed during neural crest cell delamination, which exhibits characteristics of EMT [91].

Next, we tested the hypothesis that EMT- and MET-like transitions underlie transitions between mesenchymal- and epithelial-like cell states in osteoblastic cells. Partial MET and EMT result in intermediate E/M and M/E states which exhibit dual epithelial and mesenchymal characteristics (Fig 4B), necessitating the analysis of multiple markers. We first assessed canonical markers and transcription factors associated with MET and EMT (Fig 4C-G). Cluster 1 had the highest expression of the epithelial markers *esrp1* and *cdh1*, whereas clusters 3 or 5 had the highest expression of the mesenchymal markers *cdh2/11, prrx1b, twist2/3, ctnna2/b1/b2*, and *zeb1b/2a*. The expression of *twist2/3* and *prrx1b* was highest in cluster 3. These cells match the description of *twist+* osteoprogenitors in the blastema identified by Stewart et al. [18], and which were hypothesized to be undergoing EMT. The expression of *zeb1b/zeb2a* was highest in cluster 5. Consistent with the notion that *ZEB* drives a mesenchymal identity partly by repressing *ESRP* [92], we found strongest expression of the *zeb1b/zeb2a* paralogs in cluster 5, where there is weakest *esrp1* expression. These data indicate that canonical MET and EMT components are expressed in osteoblastic cells.

To assess whether Esrp was active during fin regeneration, we examined alternative splicing of known ESRP-dependent genes using VAST-TOOLS [93]. Because our method of scRNA-seq is biased toward the 3’ end of transcripts, which may miss splicing activity that occurs near the 5’ end, we analyzed bulk RNAseq datasets of fin regeneration from Lee et al., 2020, in which paired-end libraries were sequenced [94]. Analysis detected ESRP-dependent alternative splicing in *fgfr2*, namely an increase in the epithelial isoform and decrease in the mesenchymal isoform, at 1 dpa (see Figure S7). Within this same dataset, RNAseq was performed on *sp7*+ cells [94]. In this dataset, we identified 1144 alternative splicing events (see Table S5). While we did not detect the alternative splicing of *fgfr2*, of the 1144 alternative splicing events that were found, 110 were identified to be dependent on *esrp1/2* in zebrafish (see Table S6) [69]. We concluded that *esrp1* splicing activity occurs during fin regeneration.

We next assessed whether osteoblastic cells express MET and EMT gene signatures. We constructed an orthologous EMT gene signature based on a hallmark gene set from the Molecular Signatures Database [95]. For MET, we constructed an orthologous signature based on a gene set derived from human prostate and breast cancer cell lines induced to undergo MET via OVOL1/2 expression [96]. Scoring revealed that cluster 3 had the strongest scoring for the EMT gene signature, whereas cluster 1 had the strongest scoring for the MET gene signature (Fig 4C). In regard to statistical analysis, using the Wilcoxon rank sum test with Bonferonni correction, we found that the EMT scores were significantly different between nearly all cluster pairwise comparisons except between clusters 2 and 4 (see Figure S8). MET scores were significantly different between most clusters except between clusters 1 and 2, clusters 2 and 3, and clusters 4 and 5. We also identified enrichment of these signatures in mesenchymal cell subclusters (see Figure S9).

Differential gene expression analysis revealed enrichment of MET and EMT genes in distinct osteoblast subclusters (Figure 4H, Figure S10). The top EMT markers in cluster 3 were enriched in genes associated with extracellular matrix, including matrix components (*tnc*, *col6a2*, *col5a1*, *col1a2*, *col16a1*, *col6a3*, *fbln5*, *fbln2*), collagen-binding products (*tgbi*, *lum*, *dcn*), and matrix remodeling (*mmp2*). Thus, MET- and EMT-associated gene signatures are expressed in osteoblastic cells during fin regeneration.

Finally, to elucidate cell state transitions, we ordered the osteoblastic cells in pseudotime and scored the cells for MET and EMT gene signatures (Fig 4I). Early in pseudotime, the trajectory passes through cells in clusters 1 and 2 which have high scores for MET and, in addition, low scores for osteoblast differentiation (Fig 3B) and high scores for epithelial differentiation (see Figure S11). The trajectory then progresses through cells in cluster 3. At this point in pseudotime, cells have high EMT scores, but still have low scores for osteoblast differentiation and high scores for epithelial differentiation. Finally, the trajectory progresses through cells in clusters 4 and 5. These cells have high scores for osteoblast differentiation and low scores for epithelial differentiation. Mesenchymal differentiation scores remained relatively constant throughout the trajectory. Collectively, the data suggest that in trajectory 2, as osteoblastic cells progress from less differentiated osteoblasts to more differentiated osteoblasts, they express MET-associated genes prior to expressing EMT-associated genes, and become less enriched in epithelial differentiation components.

Taken together, our studies indicate that osteoblastic cells express canonical MET and EMT components, as well as MET- and EMT-associated gene signatures. Moreover, we have identified multiple candidate trajectories underlying transitions between cell states.

### Expression of *cdh11*, *esrp1, and twist2 in situ*

To place our findings into a spatiotemporal context, we performed RNA ISH for *cdh11, esrp1, and twist2*.In some cases, we also co-stained for *runx2a*. Prior to performing RNA ISH, we analyzed our sci-RNA-seq data to help assess the relative abundance of different cell populations. Notably, osteoprogenitors that expressed both *esrp1* and *cdh11* were relatively sparse, as evidenced by the fact that *esrp1*+/*cdh11*+ cells comprised 12.7% of all osteoblast2 cells in our sci-RNA-seq analyses. Cells expressing both *runx2a/cdh11* or *esrp1/twist2* comprised ~60% and 2% of all osteoblast2 cells, respectively. In general, staining for *cdh11 and twist2* was highest in cells within the mesenchymal compartment (Fig 5DE), whereas staining for *esrp1* was highest in epithelial cells, with decreasing expression in the deeper epidermis layers (intermediate and basal epidermis) (Fig 5A-C).

**Figure 5.**
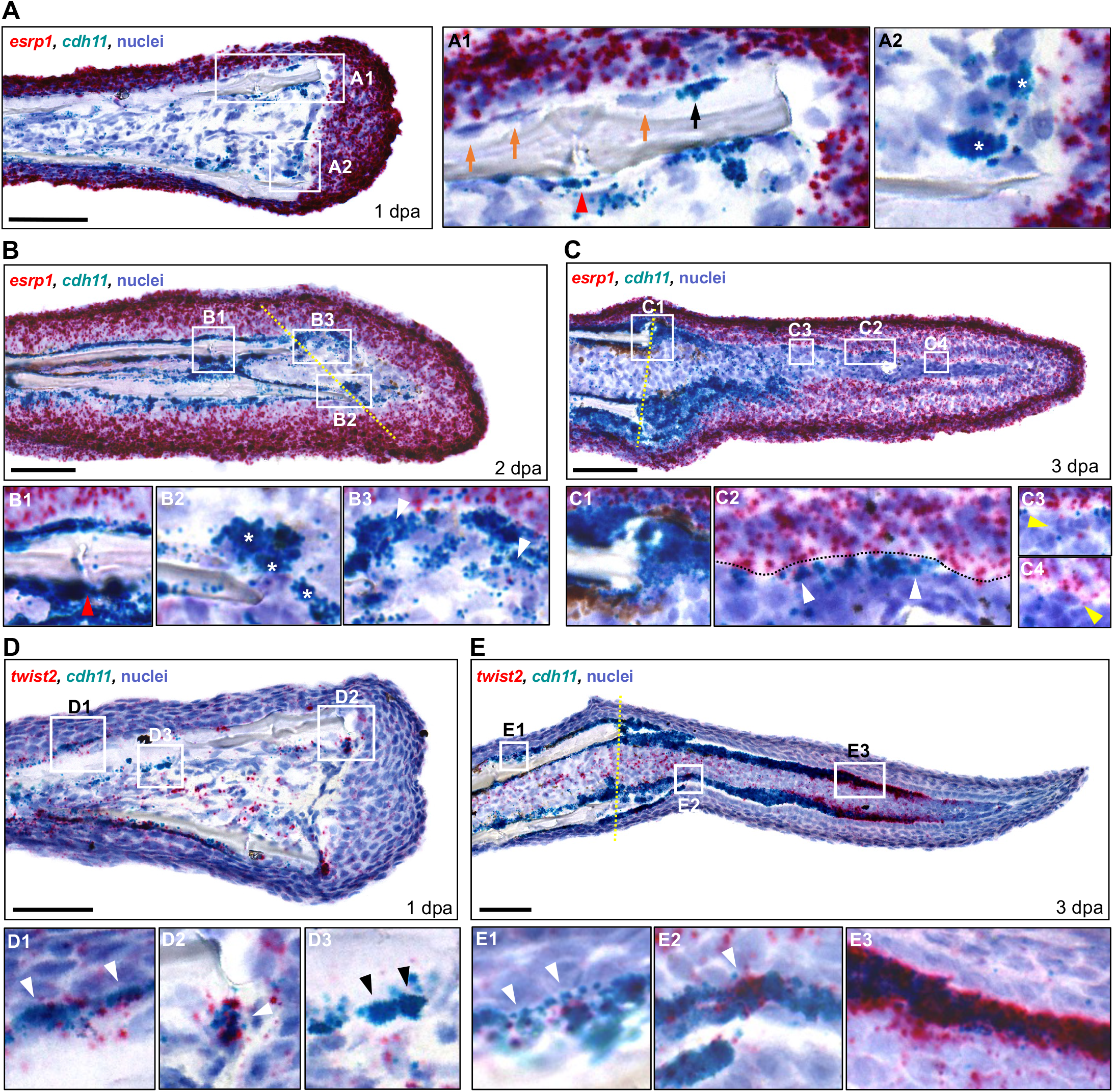
Osteoprogenitors express *esrp1* and *cdh11* in a temporal pattern during fin regeneration. (A-C) Dual RNA ISH hybridization on cryosections from 1, 2, and 3 dpa fin regenerates using RNAScope. All scale bars are 60 μm. Amputation plane is indicated with dashed yellow line. (A) Expression of *esrp1* and *cdh11* in a 1 dpa fin cryosection. (A1 inset) *cdh11* only stained some cells lining the bone (black arrow, positive staining; orange arrows, no staining). *cdh11* staining was also observed in cells around the fin ray joints (red arrowhead). (A2 inset) Cells staining for *cdh11* were observed along the amputation plane, lining the distal border of the mesenchymal compartment (stars). (B) Expression of *esrp1* and *cdh11* in a 2 dpa fin cryosection. (B1 inset) *cdh11*+ cells were prominent along the fin bone, including near the fin joint (red arrowhead). (B2 inset) Multiple *cdh11*+ cells were observed near the amputation stump (stars). (B3 inset) Arrowhead indicates a cell staining positive for both *esrp1* and *cdh11* in the forming blastema. (C) Expression of *esrp1* and *cdh11* in a 3 dpa fin cryosection. (C1 inset) Prominent *cdh11* expression is observed at the amputation stump. (C2 inset) Multiple cells showed staining for both *cdh11* and *esrp1*(arrowheads). (C3, C4 insets) Multiple cells in the lateral mesenchymal compartment were positive for *esrp1* but not *cdh11* (yellow arrowheads). (D-E) Dual RNA ISH hybridization for *twist2* and *cdh11* on cryosections from 1 and 3 dpa fin regenerates. *twist2*+/*cdh11*− cells were scattered throughout the mesenchyme at both timepoints. (D) *twist2* and *cdh11* expression in a 1 dpa fin cryosection. *twist2+/cdh11+* cells were detected along the native bone (D1 inset) and the amputation stump (D2 inset) (white arrowheads). (D3 inset) Some *cdh11+* cells along the native bone did not express *twist2* (black arrowheads). (E) *twist2* and *cdh11* expression in a 3 dpa fin cryosection. (E1 inset) *twist2*+/*cdh11*+ cells were detected along the native bone (white arrowheads). (E2 inset) Expression of *twist2* was less prominent in the proximal lateral mesenchymal compartment than *cdh11*, although small clusters of *cdh11*+ cells stained strongly for *twist2* (white arrowhead). (E3 inset) Robust *twist2* expression was observed in distal *cdh11+* cells.

We first performed dual RNA ISH for *cdh11* and *esrp1* across several regenerative stages. We first examined fins from 1 dpa (Fig 5A), a timepoint at which a wound epithelium is present but a blastema is not visible. Prior studies by Knopf et al., 2011, have shown that at this timepoint, osteoblastic dedifferentiation and migration has initiated [2]. Staining for *cdh11* was observed in bone lining cells near the amputation stump (Fig 5A1, black arrow), with decreased staining in bone lining cells in more proximal locations (Fig 5A1 orange arrows). We also observed staining for *cdh11* in cells near the fin ray joints (Fig 5A1, red arrowhead), a location known to harbor joint-derived resident osteoprogenitors [6]. Finally, staining for *cdh11* was observed in cells aligned with the amputation plane, some of which exhibited a rounded morphology (Fig 5A2, stars). Staining for *esrp1* in the mesenchymal compartment was observed occasionally in mesenchymal cells, some of which were *cdh11*+ (see Figure S12C, S13). Thus, at 1 dpa, staining for *cdh11* was present in cells located at sites that harbor dedifferentiated osteoblasts and resident osteoprogenitors, whereas staining for *esrp1* was mostly absent from these cells.

At 2 dpa, a forming blastema is visible. At this timepoint, staining for *cdh11* was noticeably increased from 1 dpa (Fig 5B). We observed increased staining in cells near the amputation stump (Fig 5B2, stars) as well as in cells near the fin ray joints (Fig 5B1, red arrowhead). Migration of osteoblasts into the blastema has been shown to occur at least between 18 hpa through 2 dpa [2]. Thus, it is possible that the broadened expression of *cdh11* in the stump from 1 to 2 dpa reflects changes in cell state associated with migration of cells during this period. Within the blastema, staining for *cdh11* was present in the lateral mesenchymal compartment (Fig 5B3). We observed staining for *esrp1* in several cells within the lateral mesenchymal compartment, and which co-stained with *cdh11* (Fig 5B3, white arrowheads). Thus, at 2 dpa, staining for *cdh11* was present in cells within the blastema that were likely to be osteoprogenitors, with staining for *esrp1* present in some of these cells.

At the 3 dpa timepoint, newly regenerated bone is visible, with more mature osteoblastic cells residing near the amputation stump, and less mature osteoprogenitors located distally [97]. Staining for *cdh11* was increased from 2 dpa (Fig 5C) and was prominent in cells lining the amputation stump (Fig 5C1), and in the lateral mesenchymal compartment where new bone was likely to form (Fig 5C2). Staining for *cdh11* generally decreased in the distal direction. In regard to *esrp1*, we observed increased staining in cells within the mesenchymal compartment compared to 2 dpa. Staining for *esrp1* was observed in cells in the lateral mesenchymal compartment in both proximal (Fig 5C3) and distal (Fig 5C4) locations. Within these cells, some co-staining of *esrp1* and *cdh11* was observed (Fig 5C2, white arrowheads). Notably, we did not observe a clear compartment of *esrp1*+ mesenchymal cells. Rather, *esrp1*+ cells were relatively sparse, and intercalated among or near *cdh11*+ cells in both proximal and distal locations within the blastema. Similar findings were observed when performing dual RNA ISH for *esrp1* and *runx2* (see Figure S12A). Thus, similar to 2 dpa, at 3 dpa, staining for *cdh11* was present in cells likely to be osteoprogenitors, with some of these cells staining for *esrp1*.

We also performed dual RNA ISH for *cdh11* and *twist2. twist2* is a known biomarker of the mesenchymal lineage and EMT, and plays a role in regulating osteoblast maturation in mammals [98]. In zebrafish, it has previously been shown to be expressed in regenerating fin at 1 dpa in the stump with whole mount ISH, and in distal *runx2+* cells at 3 dpa in longitudinal sections of regenerating fin [18]. This suggests that *twist2* expression is activated early during fin regeneration when osteoblasts are dedifferentiating (1 dpa), and is expressed in dedifferentiated, immature osteoprogenitors at 3 dpa. To determine if *cdh11* marks specific differentiation states in osteoblasts, we performed dual RNA ISH for *cdh11* and *twist2* at both timepoints. At 1 dpa, staining for *twist2* was observed in cells in the mesenchymal compartment (Figure 5D), in elongated *cdh11*+ bone lining cells along the native bone (Figure 5D1), and in cells at the amputation stump, some of which were *cdh11+* (Figure 5D2). Several *cdh11*+/*twist2*-cells were also detected along the native bone (Figure 5D3, black arrowheads). At 3 dpa (Figure 5E), we detected *cdh11+/twist2+* cells lining the native bone (Figure 5E1), and in the blastema along the lateral edges of the mesenchymal compartment, with the most prominent dual staining at the distal end where less mature osteoprogenitors are thought to reside (Figure 5E3). Small clusters of *cdh11+* cells with robust *twist2* staining were also found in the proximal blastema near the regenerating bone (Figure 5E2), where maturing osteoprogenitors are likely to reside. *cdh11*−/*twist2*+ cells were also observed in scattered cells throughout the mesenchymal compartment of both the blastema and the stump. Our findings suggest that *cdh11* is expressed in *twist2* dedifferentiating osteoblasts at 1 dpa, dedifferentiated *twist2*+ osteoprogenitors in the distal regenerate at 3 dpa, as well as more mature, *twist2*-osteoblastic cells in the proximal regenerate.

## DISCUSSION

This study provides a single cell resource for the study of osteoblastic cells during zebrafish fin regeneration, and support the contribution of MET- and EMT-associated components to this process. Our studies complement previous studies by Stewart et al., 2014, who noted dual mesenchymal and epithelial characteristics in osteoblastic cells, and first hypothesized this was due to EMT [18], by demonstrating the expression of MET and EMT signatures in osteoblastic cells. Our studies also extend the identification by Stewart et al. of *twist2*+ cells in the amputation stump and in the regenerate by showing that these cells co-express the EMT marker *cdh11*, and that the latter are enriched with EMT signatures. *In silico* evidence of a sequential EMT and MET process occurring during axolotl limb regeneration was recently reported [11]. In contrast, in their review Wong and Whited state that no EMT signatures were identified in two axolotl limb regeneration scRNA-seq studies [10, 12, 13], however methods for analysis were not provided. In this context, it is unclear whether the failure to identify EMT signatures during salamander limb regeneration in previous scRNA-seq studies reflect biological differences with zebrafish fin regeneration, or technical differences (e.g., the use of signatures different from those used here). To clarify this, it would be useful to systematically test for the same MET/EMT signatures in different sc-RNAseq datasets from different models of appendage regeneration.

In our sci-RNA-seq analyses we identified two potential trajectories underlying the topology of osteoblastic subclusters, raising questions related to potential reasons for their existence. Whereas trajectory 1 was consistent with osteoblastic differentiation from proliferative cells to form new bone, trajectory 2 appended additional cells to trajectory 1 that were intermediate between *sp7*-expressing osteoblasts (cluster 5) and proliferative cells (cluster 3). Such intermediate states might reflect cells that are in a dedifferentiating state. While substantial osteoblastic dedifferentiation occurs by 1 dpa [2], the presence of dedifferentiating cells at a later stage of regeneration (3-5 dpa) could be due to several reasons. In lineage tracing studies of zebrafish fin regeneration, it has been observed that the expansion of clones which have migrated into the blastema was temporally variable, with expansion of some clones starting as late as ~10–14 dpa [99]. In this context, it is conceivable that trajectory 2 included dedifferentiating cells which are quiescent or have not yet divided. Alternatively, as we discuss later, it is possible that there is periodic switching between de- and redifferentiation and proliferation, which might help maintain a pool of osteoblastic cells during regeneration.

It is possible that *cdh11* has distinct functions in EMT and osteoblast differentiation during fin regeneration. With respect to its potential role in EMT in fin regeneration, our study found evidence that *cdh11* is expressed in osteoblastic cells that either scored strongly for the EMT signature or hypothesized to be undergoing EMT. For example, in the 3 dpa fin regenerate, ISH revealed that *cdh11* expression in the distal osteoprogenitors was high, albeit not as high as in more proximal osteoprogenitors (Fig 6, top). These distal osteoprogenitors also stained for *twist2* and *runx2a* and likely corresponded with cells in cluster 3, which scored highest for the EMT signature. In addition, at 1 dpa, *cdh11* staining was observed in stump cells that were likely to be dedifferentiating osteoblasts, which have been speculated to be undergoing EMT [18]. It is therefore possible that *cdh11* has a function in EMT in osteoblasts. Because *cdh11* expression was elevated even further in proximal cells compared to distal osteoprogenitors, it is possible that *cdh11* may have a role in osteoblast differentiation independent of EMT during fin regeneration. More specifically, at 3 dpa, the strongest *cdh11* ISH staining was observed in the proximal osteoprogenitors, which are undergoing osteoblast differentiation and maturation, and staining was comparably reduced in the distal osteoprogenitors, which are less differentiated. From our clustering analysis, the proximal osteoprogenitors likely correspond to cluster 5 cells which not only had the strongest *cdh11* expression levels but also scored highest for osteoblastic differentiation, and differentially expressed *sp7*, a marker of osteoblast maturation. In support of a role for *cdh11* in osteoblast differentiation independent of a role in EMT, it has been previously shown that *Cdh11* is highly expressed in mesenchymal stem cells in mice during osteogenic differentiation [100]. Taken together, our study suggests that during fin regeneration, *cdh11* could play multiple roles related to EMT and osteoblast differentiation in the regenerate.

**Figure 6.**
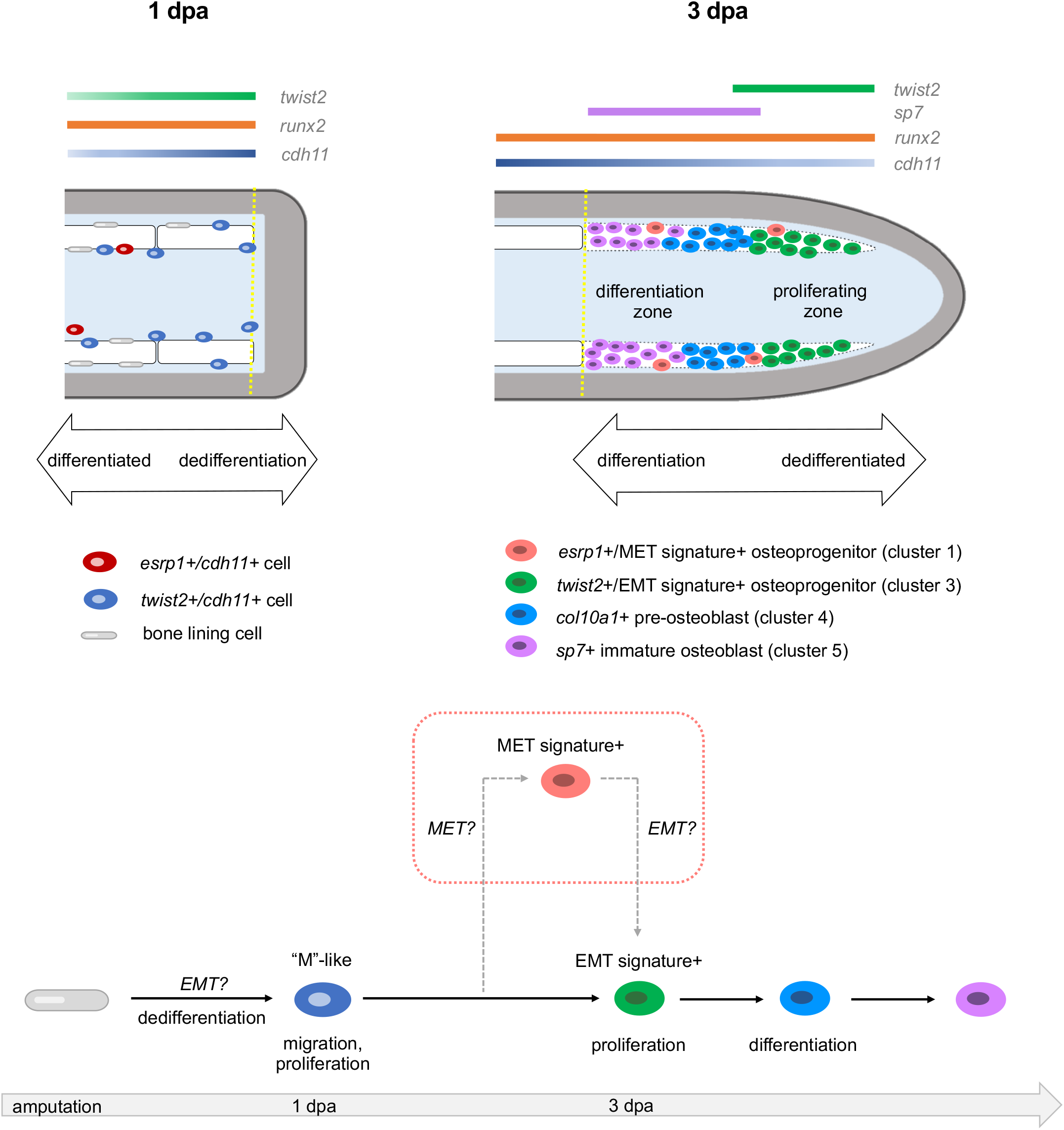
Schematics showing predicted spatial locations of the different osteoprogenitors from osteoblast2 within the fin (top) and predicted model of osteoblastic de- and redifferentiation during fin regeneration (bottom). Amputation plane is indicated with a dashed yellow line. (top, left) At 1 dpa, dedifferentiated osteoblasts expressing *cdh11* and *twist2* are observed along the native bone, at fin ray joints, and at the amputation stump. Elongated bone lining cells are still present but are *twist2*−/*cdh11*−. *esrp1*+/*cdh11*+ cells are occasionally found near the native bone. (top, right) At 3 dpa, osteoprogenitors in the blastema are located along the lateral edges of the mesenchymal compartment as indicated by the dashed white region. Early cycling osteoprogenitors expressing a strong EMT gene signature are distally located. These cells also exhibit robust expression of *runx2a* and *twist2*, and comparably less *cdh11* than in proximal regions of the regenerate. Proximal osteoprogenitors exhibit strong *cdh11* expression and have begun maturing into differentiated osteoblasts, first expressing *col10a1*, then *sp7*. Non-cycling osteoprogenitors, some of which express a strong MET gene signature, are sporadically located amongst both the proximal and distal osteoprogenitors. (bottom) Model of osteoblastic de- and redifferentiation during fin regeneration. Bone lining cells dedifferentiate into *twist2*+/*cdh11*+ cells, which are mesenchymal-like (“M”-like). These cells then migrate into the blastema and proliferate. A majority of the cells then follow the solid black arrows and progress through redifferentiation and become mature osteoblasts to regenerate bone. Some of the cells follow the dashed gray arrows, first expressing MET components before expressing EMT components, prior to differentiating into mature osteoblasts.

Previous studies have identified spatial domains in the regenerate in which cells differentially express phenotypes linked to MET and EMT, thus it is useful to consider how these domains relate to expression of MET and EMT components in our study. Stewart et al. showed that during zebrafish fin regeneration, osteoblastic cells exhibit a change in shape from more rounded cells in the regenerate to more flattened cells in proximal unamputated tissue [18]. Moreover, Mateus et al., 2015, showed that mesenchymal cells exhibit a change in shape from more round cells in the distal regenerate to more flat cells in the proximal regenerate [101]. In our studies, both *cdh11* and *twist2* were expressed in osteoblastic cells in the distal regenerate; in the proximal regenerate, *cdh11* expression was increased in osteoblastic cells, whereas *twist2* was decreased but with intermittent pockets of expression. This suggests that dual expression of *cdh11* and *twist2* is associated with a more rounded cell shape in distal cells. Future studies that correlate expression of EMT and MET components with cell shape and F-actin localization on a cell-by-cell basis could shed light on the relationship between the expression of these components and cell phenotypes linked to EMT and MET.

Our study identifies relationships between osteoblast differentiation in the regenerate and the expression of genes associated with mesenchymal/epithelial differentiation and MET/EMT. Of the osteoblast2 subclusters, cells with the lowest osteoblast differentiation scores (clusters 1 and 2) had the highest epithelial differentiation and MET scores, cells with intermediate osteoblast differentiation scores (cluster 3) had the highest EMT score and elevated mesenchymal and epithelial differentiation scores, and cells with the highest osteoblast differentiation scores (cluster 4 and 5) exhibited the lowest epithelial differentiation and MET scores. One interpretation of these data is that osteoblasts in a more epithelial-like state are less differentiated. However, this would be opposite to the findings of Stewart et al. in which epithelial characteristics were associated with the differentiated state [18]. An alternate interpretation is that osteoblasts in a less differentiated state are in a more hybrid E/M state, whereas more differentiated osteoblasts are in a less hybrid E/M state. This would be similar to what has been shown in cancer where partial EMT can enhance cancer cell stemness, and cells reversibly switch between multiple metastable hybrid E/M states resulting in a spectrum of phenotypes [9]. In this context, it is possible that the sporadic appearance of osteoblasts expressing MET signatures amongst cells that express EMT signatures reflects metastability and sporadic switching between hybrid states that are shifted toward either an epithelial or mesenchymal state, respectively. In addition, there is evidence that cancer cells in an epithelial-shifted hybrid state can exhibit a greater proliferative capacity than more mesenchymal variants [102]. Accordingly, osteoblasts in an epithelial-shifted hybrid state may be more proliferative, and as fin regeneration takes ~14-21 days to reach completion, osteoblastic cells undergoing periodic switching between epithelial- and mesenchymal-shifted hybrid states could support a replenishing pool of dedifferentiated osteoprogenitors throughout the course of regeneration.

Our study also identifies potential relationships between the processes of EMT, MET, and osteoblast re- and dedifferentiation. Within the osteoblast2 population of the regenerate, the subcluster 3 cells were marked by *twist2* expression and high EMT signature scores. These cells also had the highest proportion of cycling cells and a low osteoblast differentiation score. Thus, our data suggests that expression of EMT components is associated with the transition between cell cycling and early osteoblast differentiation and thus it is possible that it facilitates this process. Alternatively, it is possible that the expression of EMT signatures in osteoblasts at this stage is a remnant of an earlier EMT-like event. Previously, Stewart hypothesized that following amputation, osteoblasts dedifferentiate by acquiring a mesenchymal-like cell state through a process similar to EMT. Supporting this notion, at 1 dpa we observed *cdh11* and *twist2* co-expression specifically in cells that are likely dedifferentiating osteoblasts. In this context, it is plausible that while osteoblast dedifferentiation in the stump involves *cdh11*, *twist2*, and EMT signatures, expression of these components in distal, redifferentiating osteoprogenitors of the regenerate at a later timepoint reflects residual carryover from earlier processes. As for the relationship between MET and re- and de-differentiation, as we described previously, the cells in subcluster 1, which express *esrp1* and MET signatures, may represent osteoblasts that are in a dedifferentiating state at 3 and 5 dpa, timepoints that are later than when a majority of dedifferentiation occurs. Thus, it is conceivable that cells that have already expressed EMT signatures during the initial dedifferentiation may subsequently express MET signatures, followed by another round of EMT, i.e. a process of periodic de-differentiation, proliferation, and redifferentiation, which could help maintain a pool of osteoblastic progenitor cells during regeneration.

We used our data to develop a model of osteoblastic de- and redifferentiation, which we hypothesize involves two trajectories (Fig 6). The first trajectory is based on trajectory 1 and excludes cluster 1 cells that express MET gene signatures. In this trajectory (solid arrows in Fig 6), following fin amputation, bone lining cells undergo dedifferentiation and express *cdh11* and *twist2*. These *cdh11*+/*twist2*+ osteoprogenitors delaminate and migrate into the blastema, a behavior similar to epithelial cells undergoing EMT (denoted by “EMT?” in Fig 6). There, they continue to proliferate and express EMT-associated gene signatures and begin to redifferentiate into osteoblasts. Thus, in this trajectory, osteoblasts acquire a mesenchymal-like (“M”-like) cell state through a process that employs programs similar to EMT, as proposed by Stewart et al [18].

The second trajectory is based on trajectory 2 but *includes* cluster 1 cells that express MET gene signatures (dotted rectangle in Fig 6). This trajectory begins similarly as the first trajectory. However, after initially undergoing EMT, a subset of *cdh11*+ osteoprogenitors express *esrp1* and MET gene signatures. Because *esrp1*+/*cdh11*+ osteoprogenitors were relatively sparse, we hypothesize that the “EMT only” trajectory is predominant, whereas the “EMT and MET” trajectory serves as a secondary path to generate specialized cells within the blastema or to help maintain a pool of progenitors. Based on our trajectory analysis, it is conceivable that these cells undergo another round of EMT (denoted by “EMT?” in the dotted rectangle in Fig 6), upon which the cells exhibit an EMT signature and begin to redifferentiate into osteoblasts.

By identifying EMT and MET components expressed during fin regeneration, our studies provide a useful lens to identify pathways underlying de- and redifferentiation. In our studies, paralogs for *zeb1/2* and *twist* (but not *snail*) were markedly expressed in osteoblastic cells. Moreover, we observed *twist2*+/*cdh11*+ cells near the amputation stump at 1 dpa. Thus, osteoblastic dedifferentiation may involve expression of *twist*, *zeb1/2*, and *cdh11*, a combination that has been observed in certain EMT-like processes [91, 103]. With respect to the identification of MET signatures in osteoblasts, it has been hypothesized that in cancer, MET may be required for metastasis as mesenchymally-shifted cancer cells often exhibit reduced proliferative capacity. Cancer cells that have undergone a *ZEB1/2-induced* EMT are particularly associated with decreased proliferation and may as a consequence require MET to proliferate and metastasize [102]. Further studies are needed to determine whether osteoblastic cells exhibit reversibility between metastable epithelial-shifted and mesenchymal-shifted hybrid states, reminiscent of that associated with increased cancer cell stemness [9].

A growing body of studies suggests that EMT or MET pathways may be harnessed to engineer or enhance dedifferentiation in mammalian cells. For instance, early stages of induced pluripotent stem cell (iPSC) reprogramming from mouse embryonic fibroblasts require MET and transition to an epithelial-like morphology [104]. *Esrp1* has been shown to enhance reprogramming of fibroblasts to iPSCs, in part by alternative splicing of the transcription factor *Grhl1* (grainyhead like transcription factor 1) [105]. Reprogramming efficiency is also increased by early induction of EMT [106, 107]. Moreover, increased differentiation potential of breast epithelial cells similar to mesenchymal stem cells is achieved by overexpression of Snail or Twist [107, 108]. Because reprogramming induces much more dramatic changes compared to osteoblastic dedifferentiation during zebrafish fin regeneration, it is not yet clear whether EMT/MET are required for de- and redifferentiation in the latter. Future studies are needed to further elucidate MET pathways and EMT components expressed during zebrafish fin regeneration, and whether such components are necessary and sufficient for osteoblastic de- and redifferentiation in this process.

## LIMITATIONS OF THE STUDY

Some limitations of our study should be considered. First, gene expression alone is insufficient to indicate that MET or EMT is occurring [7]. In our studies, osteoblastic cells expressed several EMT-TFs (including orthologs for ZEB and TWIST, part of a core group of transcription factors that are expressed in all instances of EMT [7]), and were enriched with EMT-associated genes. However, we did not assess cell phenotypes that define EMT such as loss of apical–basal polarity, modulation of the cytoskeleton, and cell–cell adhesive properties. Thus, whether the expression of MET- and EMT-associated components in our study are reflective of actual MET and EMT, or are distinct processes that bear resemblance to MET and EMT, should be assessed in future studies. Another limitation of our study is the focus of our sci-RNA-seq analysis on a later window of regeneration (3-5 dpa). While this prevented transcriptomic examination of earlier timepoints, our RNA ISH studies allowed us to connect our single cell dataset to early osteoprogenitor states. Finally, it should be emphasized that the trajectories constructed in this study are solely based on unbiased inference, and that lineage tracing studies are needed to test their predictions.

## ACKNOWLEDGEMENTS

The authors would like to acknowledge Dr. Ting Wang for generous sharing of scRNA-seq data, as well as members of the MSBL, the UW Regeneration Club, and Dr. Lauren Saunders for helpful discussions. Research reported in this publication was supported by the Brotman Baty Institute Seattle Single Cell Initiative, a Seed Grant from the University of Washington Department of Orthopaedics and Sports Medicine, and the National Institute of Arthritis and Musculoskeletal and Skin Diseases of the National Institutes of Health under Award Number AR066061. The content is solely the responsibility of the authors and does not necessarily represent the official views of the National Institutes of Health.

## AUTHOR CONTRIBUTIONS

Conceptualization, Methodology, and Investigation: W.J.T., C.J.W., and R.Y.K.; Formal Analysis, Data Curation, Writing – Original Draft, Visualization: W.J.T. and R.Y.K.; Writing – Review & Editing: W.J.T, C.J.W., T.O., C.H.A., and R.Y.K.; Supervision, Funding Acquisition: R.Y.K.

## MAIN TABLES AND CORRESPONDING TITLES AND LEGENDS

(N/A)

## STAR METHODS

### RESOURCE AVAILABILITY

#### Lead contact

Further information and requests for resources and reagents should be directed to and will be fulfilled by the lead contact, W. Joyce Tang

#### Materials availability

This study did not generate new unique reagents.

#### Data and code availability

- Single-cell RNA-seq data have been deposited at GEO and are publicly available as of the date of publication. Accession numbers are listed in the key resources table. This paper also analyzed existing, publicly available data, the accession numbers for which are listed in the key resources table.
- This paper does not report original code.
- Any additional information required to reanalyze the data reported in this paper is available from the lead contact upon request.

### EXPERIMENTAL MODEL AND SUBJECT DETAILS

#### Zebrafish husbandry

All studies were performed on an approved protocol in accordance with the University of Washington Institutional Animal Care and Use Committee (IACUC). Zebrafish were reared and maintained at the ISCRM Aquatic Facility under a 14:10 hour light:dark cycle at 28°C and standard conditions. Adult wildtype (AB) zebrafish were used as source material for these studies. Females and males were used because robust sexual dimorphism in the rate of adult zebrafish caudal fin regeneration has not been observed [109].

### METHOD DETAILS

#### Tissue collection

Adult wildtype (AB) zebrafish were anesthetized with 0.02% MS-222 and subjected to 50% amputation of the caudal fin and allowed to recover. At various days post amputation, regenerated fin tissues were collected and either fixed overnight at 4°C in 10% neutral-buffered formalin for cryosectioning, or immediately frozen at −80°C for sequencing. 25 fins were collected at 3 dpa for the pilot sci-RNA-seq study, and 100 fins were collected for each time point for the subsequent sci-RNA-seq study.

#### Single-cell combinatorial indexing RNA-seq

Detailed methods for single-cell combinatorial indexing RNA-seq (sci-RNA-seq) are described in [20] and are available online (http://atlas.gs.washington.edu/mouse-rna). sci-RNA-seq3 was performed as previously described, using three rounds of barcoding on intact nuclei to identify transcripts with unique barcode combinations to generate single nucleus 3’ RNA-seq data [20]. Briefly, nuclei were extracted, fixed in 4% paraformaldehyde, and frozen in 100 μl of nuclei wash buffer. Nuclei were then thawed in a 37°C water bath for 5 minutes, spun down at 500 x g for 5 minutes, and permeabilized in 400 μl of permeabilization buffer (nuclei wash buffer supplemented with 0.2% Triton X-100) for 3 minutes. Nuclei were then washed, sonicated for 12 seconds on low power mode (Diagenode), washed, and resuspended in 100 μl of nuclei wash buffer. The sci-RNA-seq3 scheme of library preparation follows Cao et al., 2019, methods for paraformaldehyde-fixed nuclei with the following modifications: 2 μl of oligo-dT primers were added to each well with 80,000 nuclei for reverse transcription, the Quick Ligation kit (New England Biolabs) was used in place of T4 ligase, and tagmentation was performed using 1/40^th^ μl per well of i7-loaded TDE1 enzyme, prepared at the University of Washington following published protocols [110]. Libraries were sequenced on Illumina NovaSeq S4 flow cells at the University of Washington Northwest Genomics Center core sequencing facility with the following cycles: read 1, 34 cycles; read 2, 100 cycles; index 1, 10 cycles; index 2, 10 cycles. Approximately 23, 42, and 63 million reads were obtained from the 3 dpa pilot, 3 dpa, and 5 dpa samples, respectively. Each sample was sequenced to an average depth of ~43 million reads, resulting in an average read depth of 6,653 reads/cell. Sequencing reads were processed as previously described, with trimmed reads being mapped to the GRCz11 zebrafish genome using STAR v.2.5.2b and GRCz11.96 (ftp://ftp.ensembl.org/pub/release-96/gtf/danio_rerio/Danio_rerio.GRCz11.96.gtf.gz).

#### Cryosectioning

Fixed tissues were washed in 1X PBS/0.05%Tween20 (PBS-T) for 3×5 minutes, then dehydrated in increasing graded concentrations of methanol and stored at −20°C for up to 6 months. Tissues were processed for embedding by first rehydrating to PBS-T in increasing graded concentrations, then cryoprotected by incubation in 15% sucrose/1X PBS for 1 hour at room temperature, followed by 20% sucrose/1X PBS for 4 hours to overnight at 4°C. Embedding was done in O.C.T. compound (Fisher Scientific) in Peel-A-Way embedding molds (Polysciences, Inc., Warrington, PA) and snap frozen in either dry ice or liquid nitrogen and stored at −80°C until ready for sectioning. ~15μm sections were made on a Leica CM1850 cryostat, collected onto charged slides and allowed to dry for several hours at room temperature before being stored at −20°C.

#### In situ hybridization with RNAscope

Custom and catalog probes were purchased from the manufacturer (ACD Bio, Newark, CA). The negative control probe was provided by ACD Bio and targeted *dapB*, a gene found only in bacteria. *In situ* hybridization was performed using the RNAScope 2.5 HD Duplex Detection kit essentially as per manufacturer instructions, with the following modifications: sections were treated with Target Retrieval solution in a steamer for 8 minutes at ~100°C, and permeabilized with Protease Plus for 15 minutes at 40°C.

#### Imaging and analysis

Slides were imaged with a 40X objective on an Aperio VERSA 200 slide scanner. All images are representative of at least two different sections. Sections for 1 and 2 dpa analysis were from a single ISH experiment, and sections for 3 dpa analysis were from three separate experiments. Per the manufacturer’s recommendation, more than one spot per ten cells in a field of view at 40X magnification was considered positive staining. We verified this criterion using a negative control (see Figure S12B).

### QUANTIFICATION AND STATISTICAL ANALYSIS

#### sci-RNA-seq analysis

A pilot dataset from 25 fins collected at 3 dpa was analyzed with Seurat v3.1 using the standard recommended workflow for quality control filtering (200 < nFeatures_RNA < 2500) to remove low quality cells or potential doublets, respectively, and data processing. Briefly, the data was normalized by scaling gene content by cell and log normalizing, followed by variable feature selection using the “vst” method. The data was then scaled and principal components were calculated using the identified variable features. Clustering was performed using the Louvain algorithm, followed by non-linear dimensional reduction with UMAP to visualize the data [111]. Unbiased clustering was performed with the top 15 principal components (PCs) and with clustering resolution set to 0.5. To assign cell types to the resulting clusters, the FindAllMarkers function was used for differential gene expression analysis to identify gene markers for each cluster (logfc.threshold = 0.25, min.pct = 0.2 or 0.25, and/or min.diff.pct = 0.25 as noted in the supplemental tables).

Using the cell type label annotations of our pilot dataset as a reference, we projected the cell type labels onto the main 3 and 5 dpa datasets (n = 100 fins for each time point) using the cell type label transfer method (FindTransfer Anchors, TransferData, AddMetaData functions). The three datasets (3 dpa pilot and larger 3 and 5 dpa datasets) were then combined using the merge function in Seurat. We did not employ Seurat integration workflows [112] since such workflows were designed for situations where substantial differences such as those arising from differences in scRNA-seq technologies or modalities need to be harmonized, which did not apply to our data. Moreover, while we did not observe evidence of batch effects in our data—which would be manifested as cells from each timepoint clustering together—the cell type label transfer workflow accounts for batch differences if present.

The standard quality control filtering and data processing workflow was repeated on the merged datasets (200 < nFeatures_RNA < 2500, number of PCs = 15, resolution = 0.5). To subcluster the osteoblast population, we subset the Osteoblast2 cells from the combined dataset and again applied the standard Seurat data processing workflow (number of PCs = 15, resolution = 0.8). Osteoblast scores for gene expression programs were calculated using the AddModuleScore function with the appropriate gene sets and using default parameters. All plots were generated in R with either Seurat, base R graphics, or with the dittoSeq package [113]. “Open” trajectory analysis was performed with Slingshot v1.2 [64] using appropriate data embeddings (either UMAP or the first two PCs from principal component analysis). “Closed” trajectory analysis was performed using the function “principal_curve” (available in base R > 3.0) and a periodic lowess smoother with appropriate data embeddings. Pseudotime values were obtained by computing projections onto the closed curve.

#### Mapping data onto existing fin regeneration data

Hou et al., 2020, generously shared the Seurat object containing their fin regeneration data. We used the FindTransferAnchors and TransferData functions in Seurat to project the tissue type annotations from our pilot 3 dpa dataset onto the Hou et al. dataset and assessed the efficacy of the mapping by confirming the expression of known marker genes in each tissue type. To determine the number of cells from the Hou et al. dataset which mapped to osteoblast1 and osteoblast2, we counted the number of cells from the Hou et al. dataset which were predicted to belong in the osteoblast1 and osteoblast2 clusters.

#### Cell Scoring, cell cycle analysis, statistical analyses

Gene sets associated with cellular functions such as epithelial differentiation, mesenchymal differentiation, MET and EMT, were obtained from various sources including the Molecular Signatures Database (MSigDB), the Gene Ontology database and the literature, to create “gene signatures” for each cellular function. Using the AddModuleScore function, cells were scored for the expression of each gene signature to calculate the difference between the average expression level of each gene signature and a set of control genes randomly picked as previously described [114]. Zebrafish orthologs for the genes in each gene set were identified if possible. Gene signatures for epithelial differentiation, mesenchymal differentiation, MET and EMT consisted of approximately 100-300 gene orthologs. To test for statistical differences of the scores between each cluster, we first tested for normal distribution using the Shapiro-Wilks test. As the data was not normally distributed, we used the non-parametric Kruskal-Wallis test followed by pairwise comparisons using the Wilcoxon rank sum test with Bonferroni correction. Cell cycle analysis was performed using the CellCycleScoring function in Seurat, which classifies each cell as being in either G2M, S, or G1 phase based on the expression of G2/M and S phase markers [114]. To test if the proportion of cycling to non-cycling cells was significantly different between clusters, we performed a difference of proportions test followed by a pairwise comparison using Bonferroni correction.

#### Alternative splicing analysis

The Vertebrate Alternative Splicing and Transcription Tools (VAST-TOOLS) software [93] was used to profile alternative splicing events in bulk RNA-seq data (GSE126701) [94]. Data from experimental replicates were merged, and differential splicing analysis was performed using the “danRer10” library of alternative splicing profiles in zebrafish as reference (downloaded from the VastDB site, https://vastdb.crg.eu/wiki/Main_Page). All comparisons were performed with the “compare” module and implemented with the following parameters: --min_dPSI 15 --min_range 5 --p_IR. A custom R script was used to compare the alternative splicing events against those that were previously annotated as *esrp-*dependent [69].

### ADDITIONAL RESOURCES

We have used the ShinyCell package in R [115] to create a website application that allows users to visually explore our datasets in various plotted formats (UMAP dimensional plots, dot plots, etc), which is available at this URL: https://msblgroup.shinyapps.io/finregeneration/.

## SUPPLEMENTAL ITEM TITLES AND LEGENDS

Figure S1. Cell populations identified at each timepoint, Related to Figure 1. (A) UMAP plots of individual single-cell RNA-seq datasets, separated by the time of tissue collection (3A = 3 dpa pilot experiment, 3B = 3 dpa, 5 = 5 dpa). The three datasets were pooled together for unsupervised clustering analysis with Seurat. Cells are colored by cell type. The plots show all 12 cell populations were present at each timepoint. (B) Same as in (A) except cells are colored by dpa. (C) Barplot showing percentage of cell types present at each timepoint.

Figure S2. Distribution and analysis of osteoblast differentiation scores for the osteoblast2 cluster, Related to Figure 3. Violin plot overlaid with a boxplot visualizing the osteoblast differentiation scores for each subcluster of the osteoblast2 population. The table at right shows the p-values resulting from the statistical analyses used to assess for differences in osteoblast differentiation scores between clusters (Wilcoxon rank sum test with Bonferroni correction).

Figure S3. Distribution and analysis of cell cycle scores for the osteoblast2 cluster, Related to Figure 3. (Top) Results from cell cycle scoring analysis performed on osteoblast2 subclusters. Values indicate number or percent of cells in G1, G2M, or S-phase of the cell cycle. (Bottom) The table below shows the p-values resulting from statistical analyses used to assess for differences in number of cells per cell cycle phase (difference of proportions test with Bonferroni correction).

Figure S4. Analysis of doublet prediction scores, Related to Figure 3. (A) Violin plot showing scores from the doublet prediction analysis performed on pooled 3 dpa pilot, 3 dpa, and 5 dpa datasets. Doublet prediction was performed using the computeDoubletDensity function in the scDblFinder package in R. (B) Violin plot visualizing scores from doublet prediction analysis performed on osteoblast2 subclusters. (C) Scatterplot showing doublet score vs *esrp1* expression in osteoblast2 cells. Cells are colored by subcluster, and the Pearson correlation coefficient between the two features is displayed at top (*r* = 0.14).

Figure S5. Enrichment analysis of GO terms and signaling pathway members for the osteoblast2 cluster, Related to Figure 3. (A) Dotplot of GO term enrichment in the osteoblast2 subclusters. (B) Dotplot showing enrichment of pathway members (EGFR, RA) and stimulation-responsive genes (MAPK, PI3K, TNF-A, Wnt, H2O2, IL1, JAK-STAT, TFG-B, VEGF, TLR) in the osteoblast2 subclusters.

Figure S6. Slingshot pseudotime analysis produces similar trajectories when performed with either UMAP or PCA embeddings, Related to Figure 3. (A) Slingshot trajectory analysis of osteoblast2 using UMAP embeddings. (B) Slingshot trajectory analysis of osteoblast2 using the first two principal components (PC1, PC2) from principal component analysis. (C) Comparison of cell orderings derived from PCA (y-axis) and UMAP (x-axis).

Figure S7. ESRP-dependent alternative splicing of *fgfr2* during fin regeneration, Related to Figure 4. Alternative splicing analysis of bulk RNA-seq transcripts from fin tissues collected at 0, 1, and 4 dpa during fin regeneration detected an increase in the epithelial *fgfr2* splice isoform (pink line, splice event ID DreEX0032683) and a decrease in the mesenchymal *fgfr2* splice isoform (cyan line, splice event ID DreEX0032684) at 1 dpa. Data was obtained from Lee et al., 2020, GSE126701. PSI = “percent spliced in”, the ratio of normalized read counts supporting exon inclusion to the total number of normalized reads supporting exon inclusion and exclusion.

Figure S8. Analysis of EMT and MET scores for the osteoblast2 cluster, Related to Figure 4. (A) Violin plot overlaid with a boxplot showing EMT scores for each subcluster of the osteoblast2 population. The table at right shows the p-values resulting from the statistical analyses used to asses for differences in EMT scores between clusters. (B) Violin plot overlaid with boxplot showing MET scores for each subcluster of the osteoblast2 population. The table at right shows the corresponding p-values resulting from the statistical analyses. A Wilcoxon rank sum test followed by pairwise comparisons with Bonferroni correction was used for both analyses.

Figure S9. Unsupervised clustering of mesenchyme1 cells shows enrichment of similar EMT and MET scores, Related to Figure 4. (A) UMAP plot of mesenchyme1 cells showing 7 clusters of cells. (B) Dotplot of mesenchyme1 clusters showing enrichment of canonical MET and EMT markers and gene signatures.

Figure S10. Clustered heatmap of MET and EMT gene expression in osteoblast2 subclusters, Related to Figure 4. Expression levels are averaged across each subcluster, and Euclidean distances were used to determine clustering relationships.

Figure S11. Analysis of differentiation scores in pseudotime-ordered osteoblast2 cells, Related to Figure 4. Pseudotime-ordered osteoblast2 cells showing scores for (A) epithelial differentiation, (B) mesenchymal differentiation, and (c) osteoblast differentiation plotted along the y-axis. Tables at right show corresponding p-values resulting from Wilcoxon rank sum test with Bonferroni correction.

Figure S12. Dual RNA in situ hybridization (ISH) in cryosections of regenerating fin tissues at various timepoints, Related to Figure 5. Yellow dashed line indicates the amputation plane. (A) Dual RNA *in situ* hybridization for *esrp1* and *runx2a* in a 3 dpa fin cryosection. Multiple cells (arrowheads in A1, A2) in both proximal and distal locations within the blastema showed costaining for *runx2a* and *esrp1*. Scale bar = 100 μm. (B) Negative control ISH in a 3 dpa fin cryosection. Both control probes target *dapB*, a gene found only in bacteria. Scale bar = 100 μm. (C) Expression of *esrp1* and *cdh11* in a 1 dpa fin cryosection. (C1 inset) *esrp1* was infrequently observed in mesenchymal cells, some of which were *cdh11*-(yellow arrowheads). (C2, C3 insets) Several *cdh11+* cells near the native bone expressed *esrp1* as well (white arrowheads). Scale bar = 60 μm.

Figure S13. Violin plots visualizing expression levels of *cdh11, esrp1*, and *twist2* in the pooled datasets, Related to Figure 5. Clusters are sorted in order of highest to lowest average expression.

Table S1. Top 30 markers for each cell cluster identified in pooled 3 and 5 dpa dataset, Related to Figure 1.

Table S2. Differential gene expression analysis comparing osteoblast1 and osteoblast2 clusters, Related to Figure 2.

Table S3. Differential gene expression analysis comparing osteoblast clusters (osteoblast1 and osteoblast2) to non-osteoblastic clusters, Related to Figure 2.

Table S4. Top 20 cluster markers for osteoblast2, Related to Figure 3.

Table S5. Alternative splicing analysis comparing osteoblasts at 0 dpa vs 4 dpa, Related to Figure 4.

Table S6. *esrp1/2*-dependent alternative splicing events in osteoblasts, Related to Figure 4.

