## Supplemental Figures S1-S13 for "Single-cell resolution of MET- and EMT-like programs in osteoblasts during zebrafish fin regeneration"

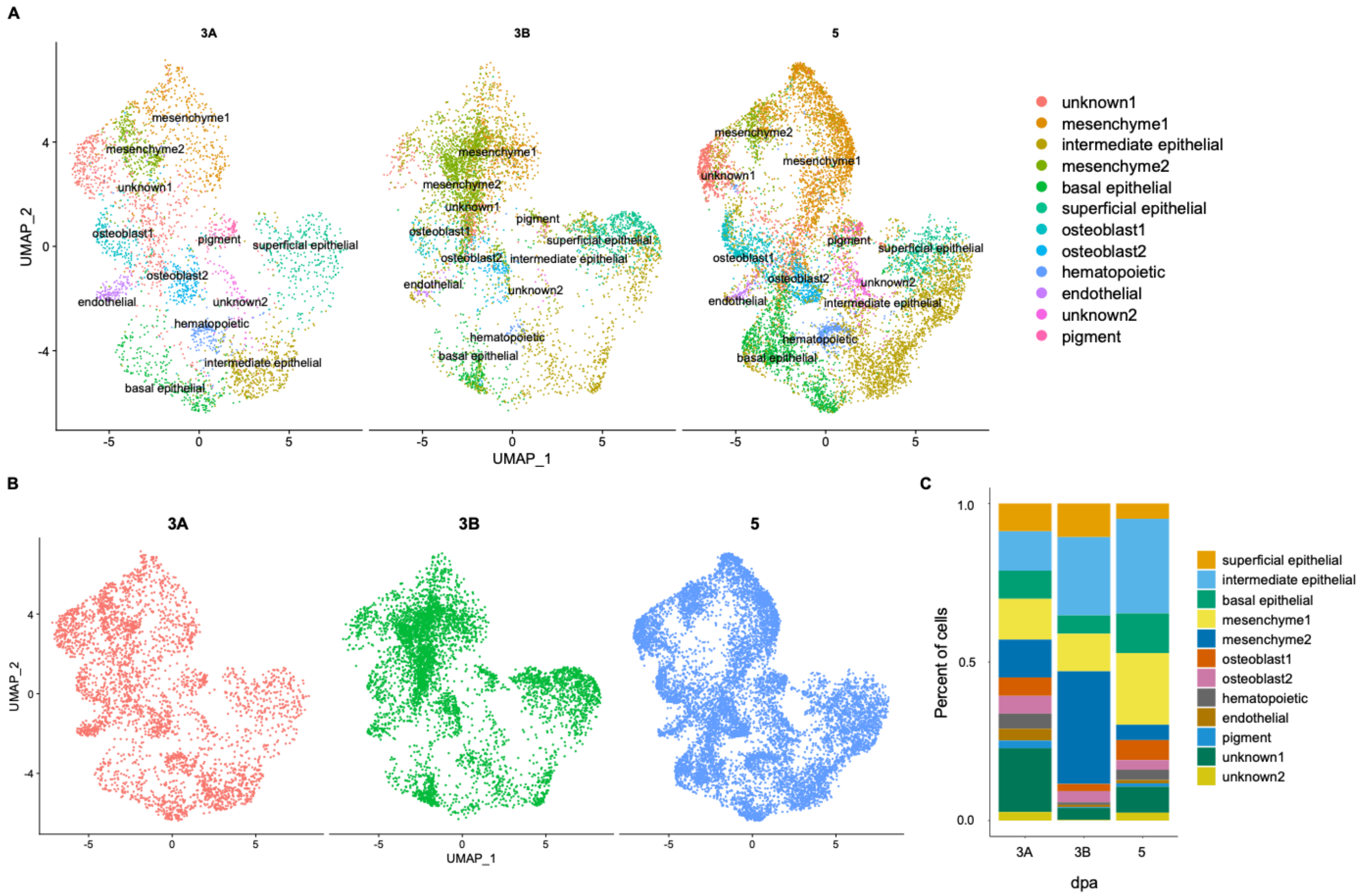


Figure S1. Cell populations identified at each timepoint, Related to Figure 1. (A) UMAP plots of individual single-cell RNA-seq datasets, separated by the time of tissue collection (3A = 3 dpa pilot experiment, 3B = 3 dpa, 5 = 5 dpa). The three datasets were pooled together for unsupervised clustering analysis with Seurat. Cells are colored by cell type. The plots show all 12 cell populations were present at each timepoint. (B) Same as in (A) except cells are colored by dpa. (C) Barplot showing percentage of cell types present at each timepoint.

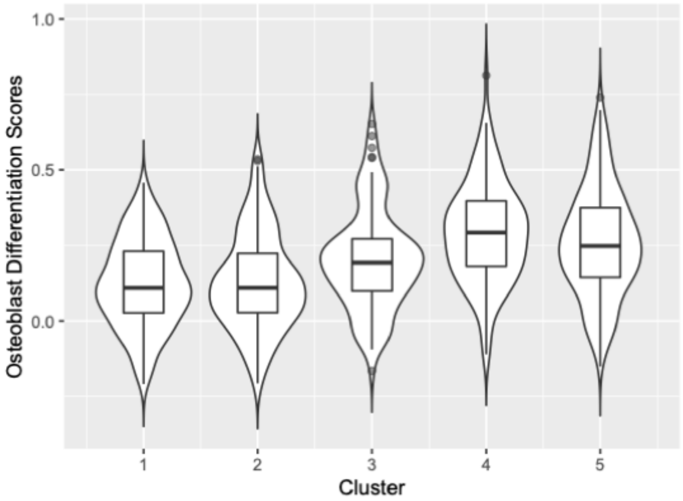


| P-values from post hoc pairwise comparisons (significant p-values in red) | | | | |
|---|---|---|---|---|
| | 1 | 2 | 3 | 4 |
| 2 | 1.00000 | - | - | - |
| 3 | 0.00218 | 0.00497 | - | - |
| 4 | 3.3e-14 | 1.1e-11 | 0.00072 | - |
| 5 | 2.4e-11 | 4.0e-09 | 0.07497 | 0.89782 |

Figure S2. Distribution and analysis of osteoblast differentiation scores for the osteoblast2 cluster, Related to Figure 3. Violin plot overlaid with a boxplot visualizing the osteoblast differentiation scores for each subcluster of the osteoblast2 population. The table at right shows the p-values resulting from the statistical analyses used to assess for differences in osteoblast differentiation scores between clusters (Wilcoxon rank sum test with Bonferroni correction).

| Number of cells in each cell cycle phase (per osteoblast2 cluster) | | | | | | | | | | |
|---|---|---|---|---|---|---|---|---|---|---|
| | G1 | G2M | S | Total | Cycling (G2M + S) | % G1 | % G2M | % S | % Cycling | % Non-cycling |
| **1** | 82 | 44 | 45 | 171 | 89 | 0.4795322 | 0.2573099 | 0.2631579 | 0.5204678 | 0.4795321 |
| **2** | 54 | 40 | 30 | 124 | 70 | 0.4354839 | 0.3225806 | 0.2419355 | 0.5645161 | 0.4354838 |
| **3** | 22 | 51 | 31 | 104 | 82 | 0.2115385 | 0.4903846 | 0.2980769 | 0.7884615 | 0.2115384 |
| **4** | 48 | 34 | 31 | 113 | 65 | 0.4247788 | 0.3008850 | 0.2743363 | 0.5752212 | 0.4247787 |
| **5** | 128 | 35 | 25 | 188 | 60 | 0.6808511 | 0.1861702 | 0.1329787 | 0.3191489 | 0.6808510 |

| P-values from post hoc pairwise comparisons (significant p-values in red) | | | | |
|---|---|---|---|---|
| | 1 | 2 | 3 | 4 |
| 2 | 1.00000 | - | - | - |
| 3 | 0.00016 | 0.00600 | - | - |
| 4 | 1.00000 | 1.00000 | 0.01320 | - |
| 5 | 0.00170 | 0.00029 | 4e-13 | 0.00022 |

Figure S3. Distribution and analysis of cell cycle scores for the osteoblast2 cluster, Related to Figure 3. (Top) Results from cell cycle scoring analysis performed on osteoblast2 subclusters. Values indicate number or percent of cells in G1, G2M, or S-phase of the cell cycle. (Bottom) The table below shows the p-values resulting from statistical analyses used to assess for differences in number of cells per cell cycle phase (difference of proportions test with Bonferroni correction).

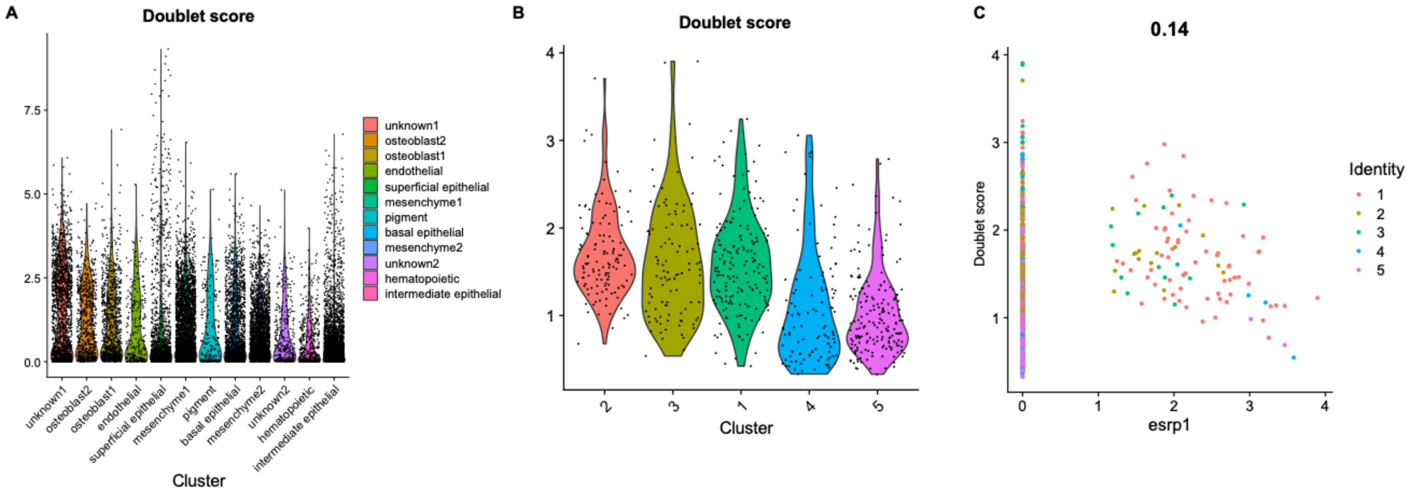


Figure S4. Analysis of doublet prediction scores, Related to Figure 3. (A) Violin plot showing scores from the doublet prediction analysis performed on pooled 3 dpa pilot, 3 dpa, and 5 dpa datasets. Doublet prediction was performed using the computeDoubletDensity function in the scDblFinder package in R. (B) Violin plot visualizing scores from doublet prediction analysis performed on osteoblast2 subclusters. (C) Scatterplot showing doublet score vs *esrp1* expression in osteoblast2 cells. Cells are colored by subcluster, and the Pearson correlation coefficient between the two features is displayed at top ($r$ = 0.14).

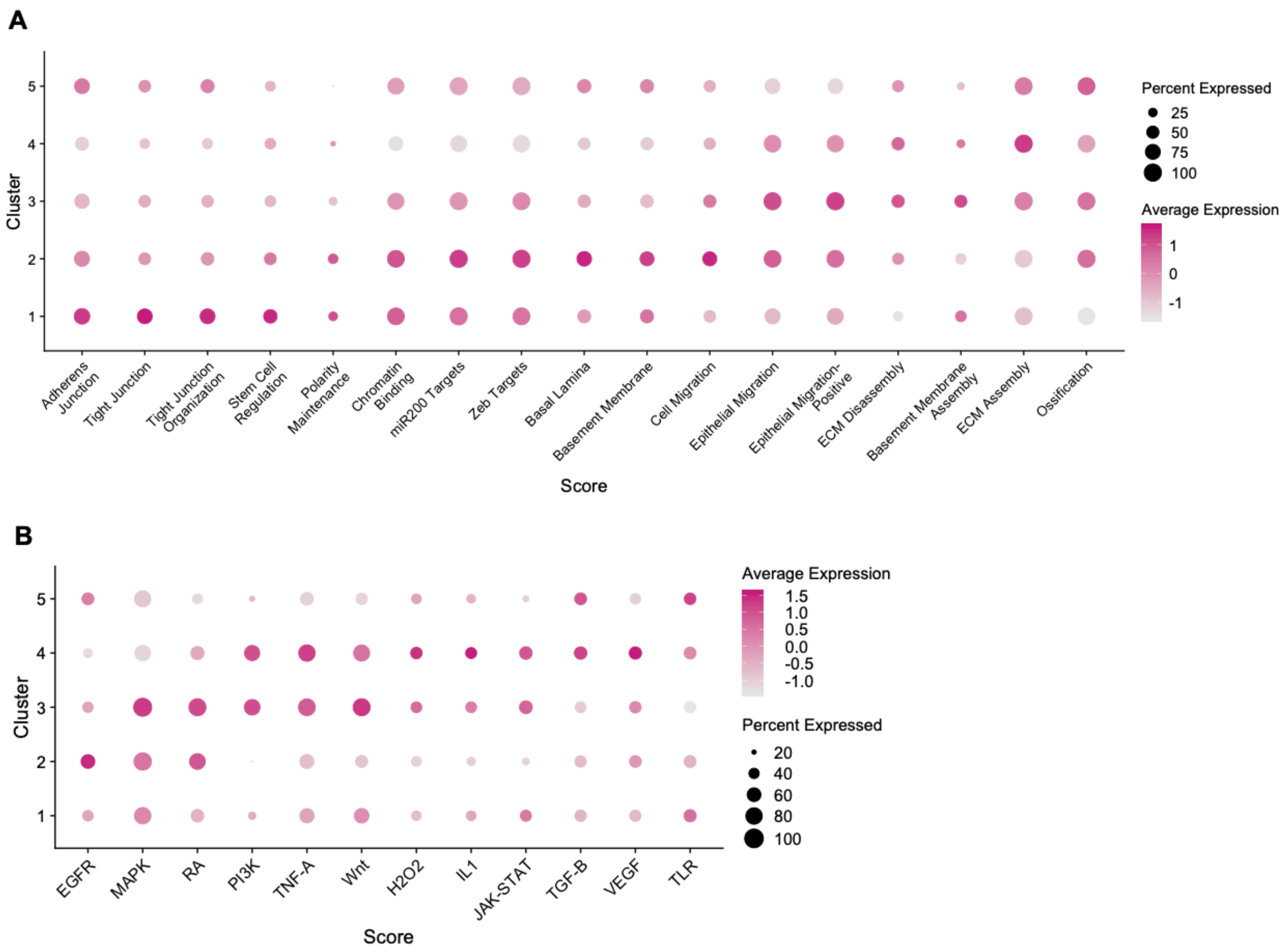


Figure S5. Enrichment analysis of GO terms and signaling pathway members for the osteoblast2 cluster, Related to Figure 3. (A) Dotplot of GO term enrichment in the osteoblast2 subclusters. (B) Dotplot showing enrichment of pathway members (EGFR, RA) and stimulation-responsive genes (MAPK, PI3K, TNF-A, Wnt, H2O2, IL1, JAK-STAT, TFG-B, VEGF, TLR) in the osteoblast2 subclusters.

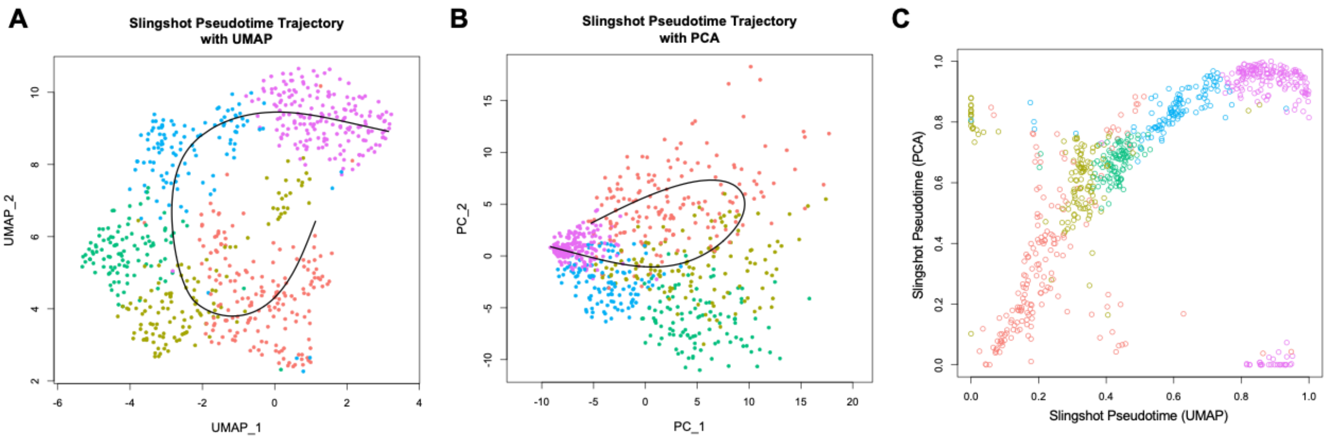


Figure S6. Slingshot pseudotime analysis produces similar trajectories when performed with either UMAP or PCA embeddings, Related to Figure 3. (A) Slingshot trajectory analysis of osteoblast2 using UMAP embeddings. (B) Slingshot trajectory analysis of osteoblast2 using the first two principal components (PC1, PC2) from principal component analysis. (C) Comparison of cell orderings derived from PCA (y-axis) and UMAP (x-axis).

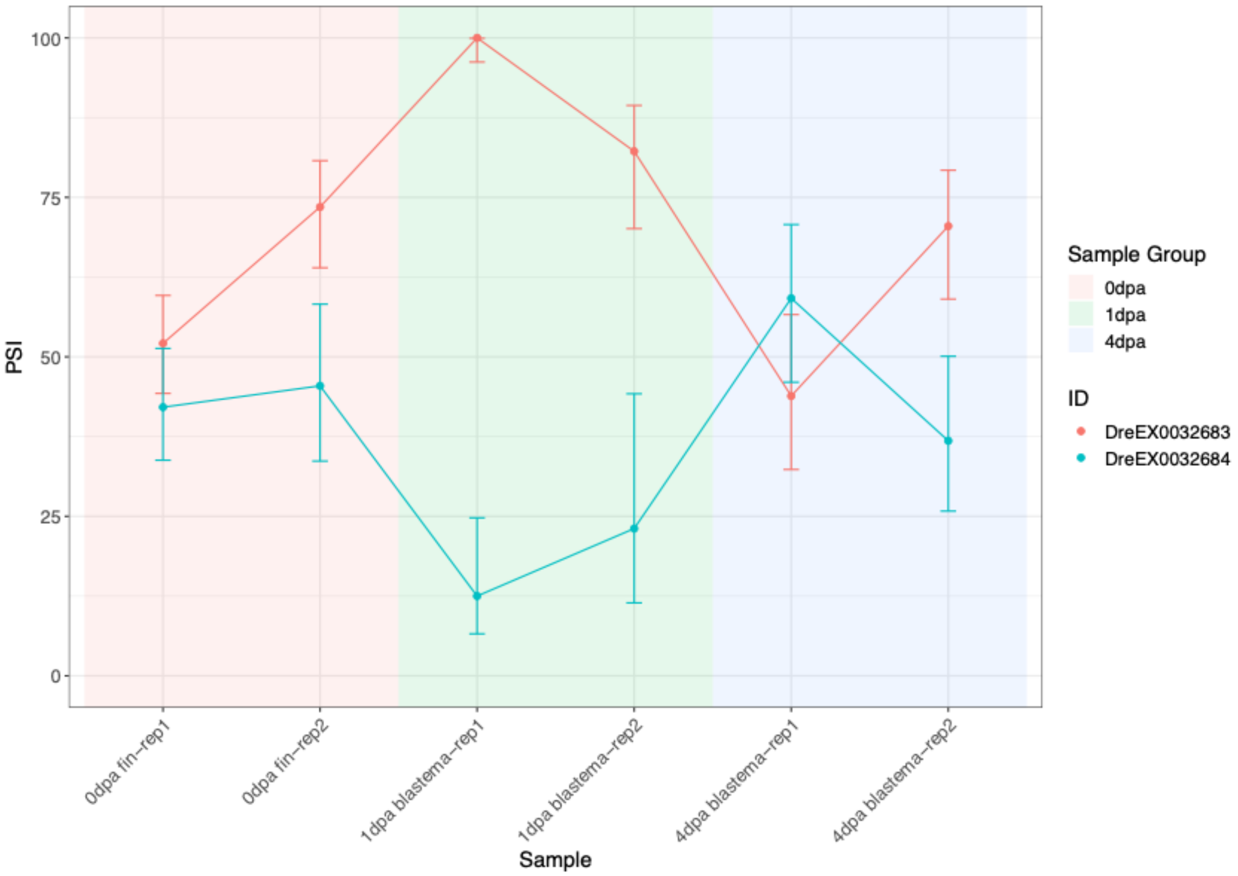


Figure S7. ESRP-dependent alternative splicing of *fgfr2* during fin regeneration, Related to Figure 4. Alternative splicing analysis of bulk RNA-seq transcripts from fin tissues collected at 0, 1, and 4 dpa during fin regeneration detected an increase in the epithelial *fgfr2* splice isoform (pink line, splice event ID DreEX0032683) and a decrease in the mesenchymal *fgfr2* splice isoform (cyan line, splice event ID DreEX0032684) at 1 dpa. Data was obtained from Lee et al., Genome Biology 2020, GSE126701. PSI = “percent spliced in”, the ratio of normalized read counts supporting exon inclusion to the total number of normalized reads supporting exon inclusion and exclusion.

**A**

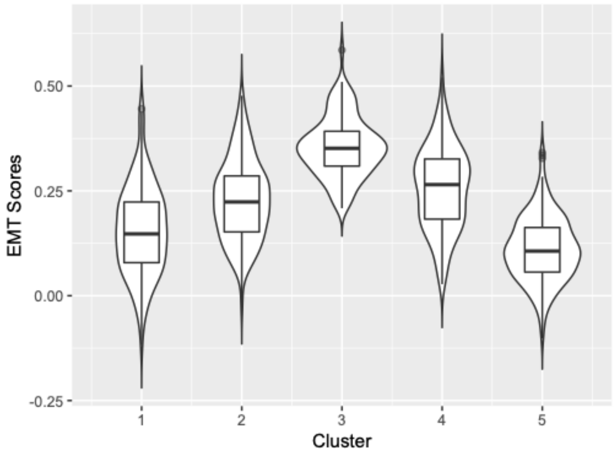


| P-values for pairwise comparisons of EMT scores (significant p-values in red) | | | | |
|---|---|---|---|---|
| | 1 | 2 | 3 | 4 |
| 2 | 1.6e-07 | - | - | - |
| 3 | < 2e-16 | < 2e-16 | - | - |
| 4 | 1.5e-12 | 0.19686 | 2.9e-12 | - |
| 5 | 0.00017 | < 2e-16 | < 2e-16 | < 2e-16 |

**B**

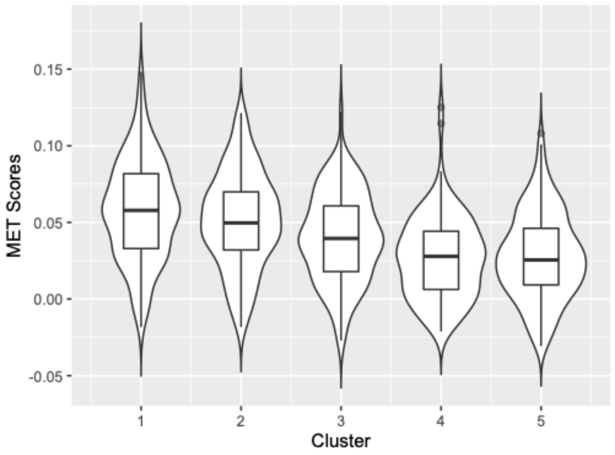


| P-values for pairwise comparisons of MET scores (significant p-values in red) | | | | |
|---|---|---|---|---|
| | 1 | 2 | 3 | 4 |
| 2 | 1.00000 | - | - | - |
| 3 | 0.00026 | 0.08348 | - | - |
| 4 | 5.6e-14 | 9.8e-09 | 0.00225 | - |
| 5 | 5.8e-16 | 2.0e-09 | 0.00331 | 1.00000 |

Figure S8. Analysis of EMT and MET scores for the osteoblast2 cluster, Related to Figure 4. (A) Violin plot overlaid with a boxplot showing EMT scores for each subcluster of the osteoblast2 population. The table at right shows the p-values resulting from the statistical analyses used to asses for differences in EMT scores between clusters. (B) Violin plot overlaid with boxplot showing MET scores for each subcluster of the osteoblast2 population. The table at right shows the corresponding p-values resulting from the statistical analyses. A Wilcoxon rank sum test followed by pairwise comparisons with Bonferroni correction was used for both analyses.

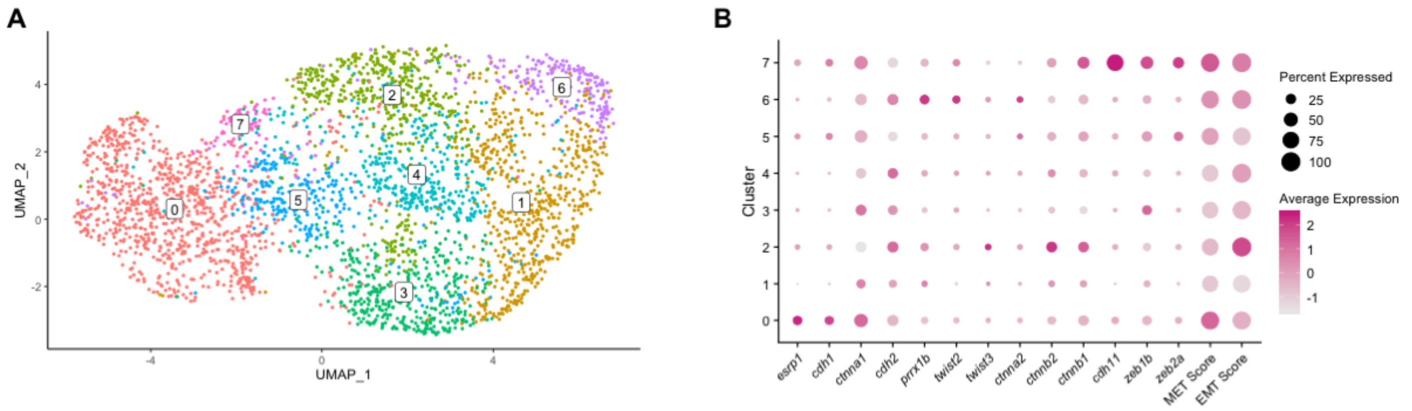


Figure S9. Unsupervised clustering of mesenchyme1 cells shows enrichment of similar EMT and MET scores, Related to Figure 4. (A) UMAP plot of mesenchyme1 cells showing 7 clusters of cells. (B) Dotplot of mesenchyme1 clusters showing enrichment of canonical MET and EMT markers and gene signatures.

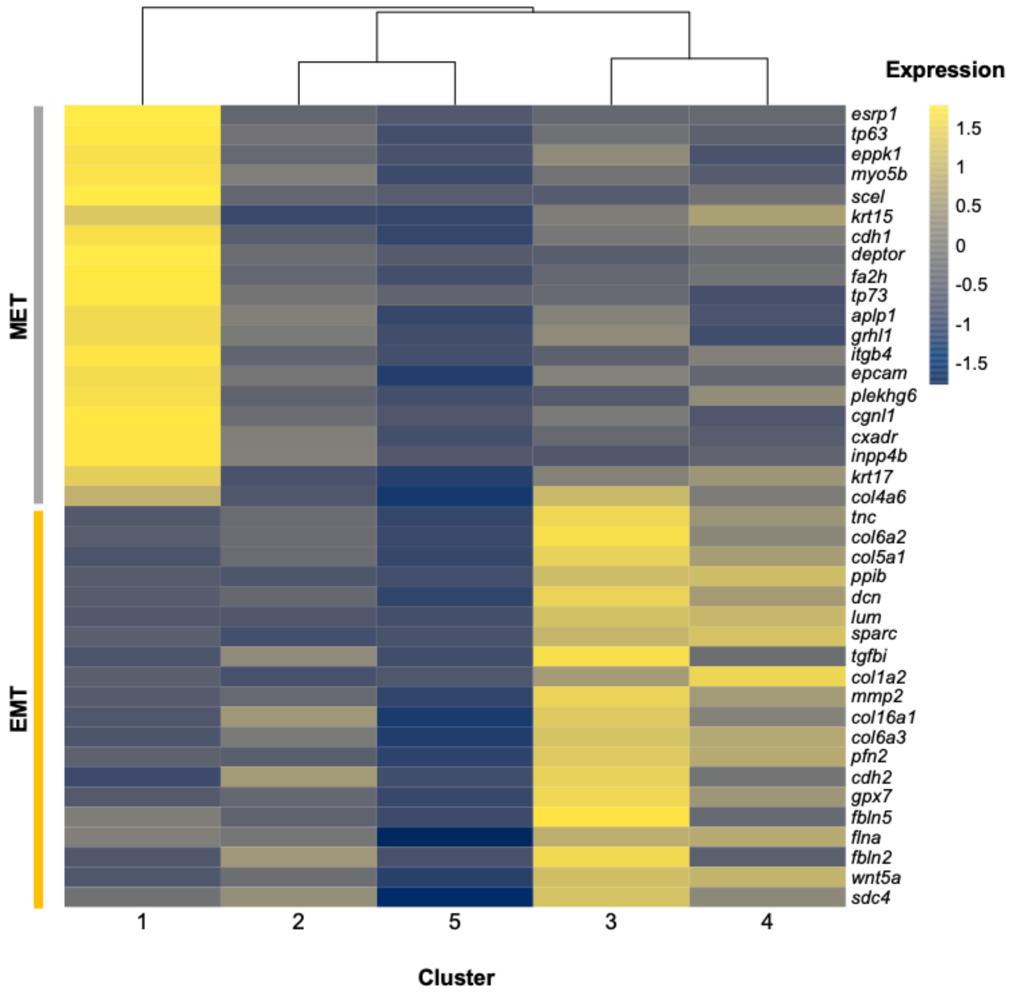


Figure S10. Clustered heatmap of MET and EMT gene expression in osteoblast2 subclusters, Related to Figure 4. Expression levels are averaged across each subcluster, and Euclidean distances were used to determine clustering relationships.

A

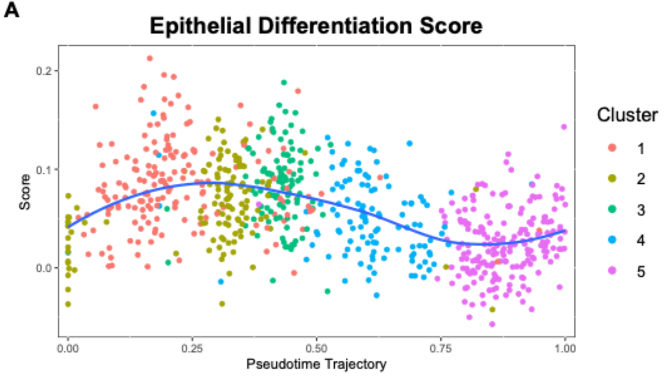


| P-values for pairwise comparisons of Epithelial Differentiation scores (significant p-values in red) | | | | |
|---|---|---|---|---|
| | 1 | 2 | 3 | 4 |
| 2 | 0.0010 | - | - | - |
| 3 | 1.0000 | 0.0015 | - | - |
| 4 | 8.7e-10 | 0.0427 | 1.1e-08 | - |
| 5 | < 2e-16 | 1.5e-14 | < 2e-16 | 6.0e-06 |

B

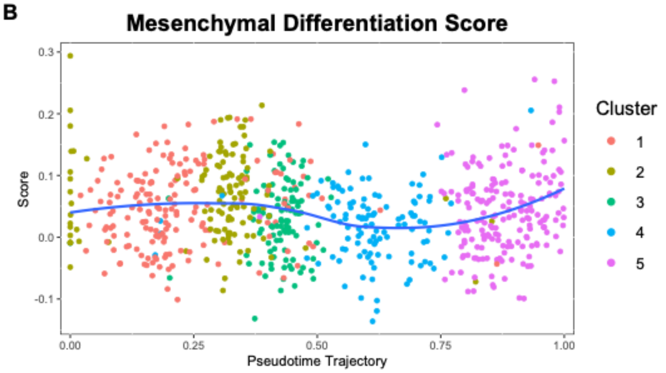


| P-values for pairwise comparisons of Mesenchymal Differentiation scores (significant p-values in red) | | | | |
|---|---|---|---|---|
| | 1 | 2 | 3 | 4 |
| 2 | 0.203 | - | - | - |
| 3 | 0.337 | 0.001 | - | - |
| 4 | 0.002 | 4.27e-07 | 1.000 | - |
| 5 | 1.000 | 0.021 | 1.000 | 0.026 |

C

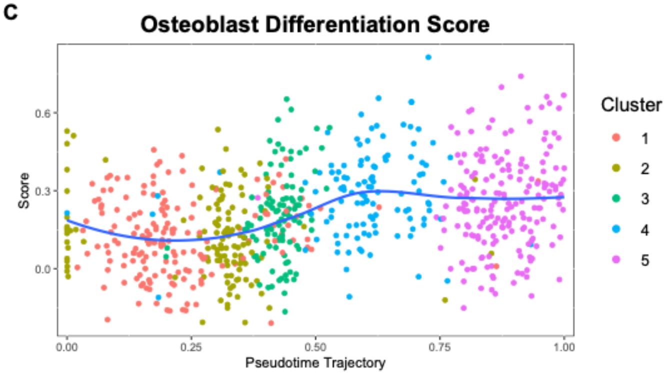


| P-values for pairwise comparisons of Osteoblast Differentiation scores (significant p-values in red) | | | | |
|---|---|---|---|---|
| | 1 | 2 | 3 | 4 |
| 2 | 1.00000 | - | - | - |
| 3 | 0.00218 | 0.00497 | - | - |
| 4 | 3.3e-14 | 1.1e-11 | 0.00072 | - |
| 5 | 2.4e-11 | 4.0e-09 | 0.07497 | 0.89782 |

Figure S11. Analysis of differentiation scores in pseudotime-ordered osteoblast2 cells, Related to Figure 4. Pseudotime-ordered osteoblast2 cells showing scores for (A) epithelial differentiation, (B) mesenchymal differentiation, and (c) osteoblast differentiation plotted along the y-axis. Tables at right show corresponding p-values resulting from Wilcoxon rank sum test with Bonferroni correction.

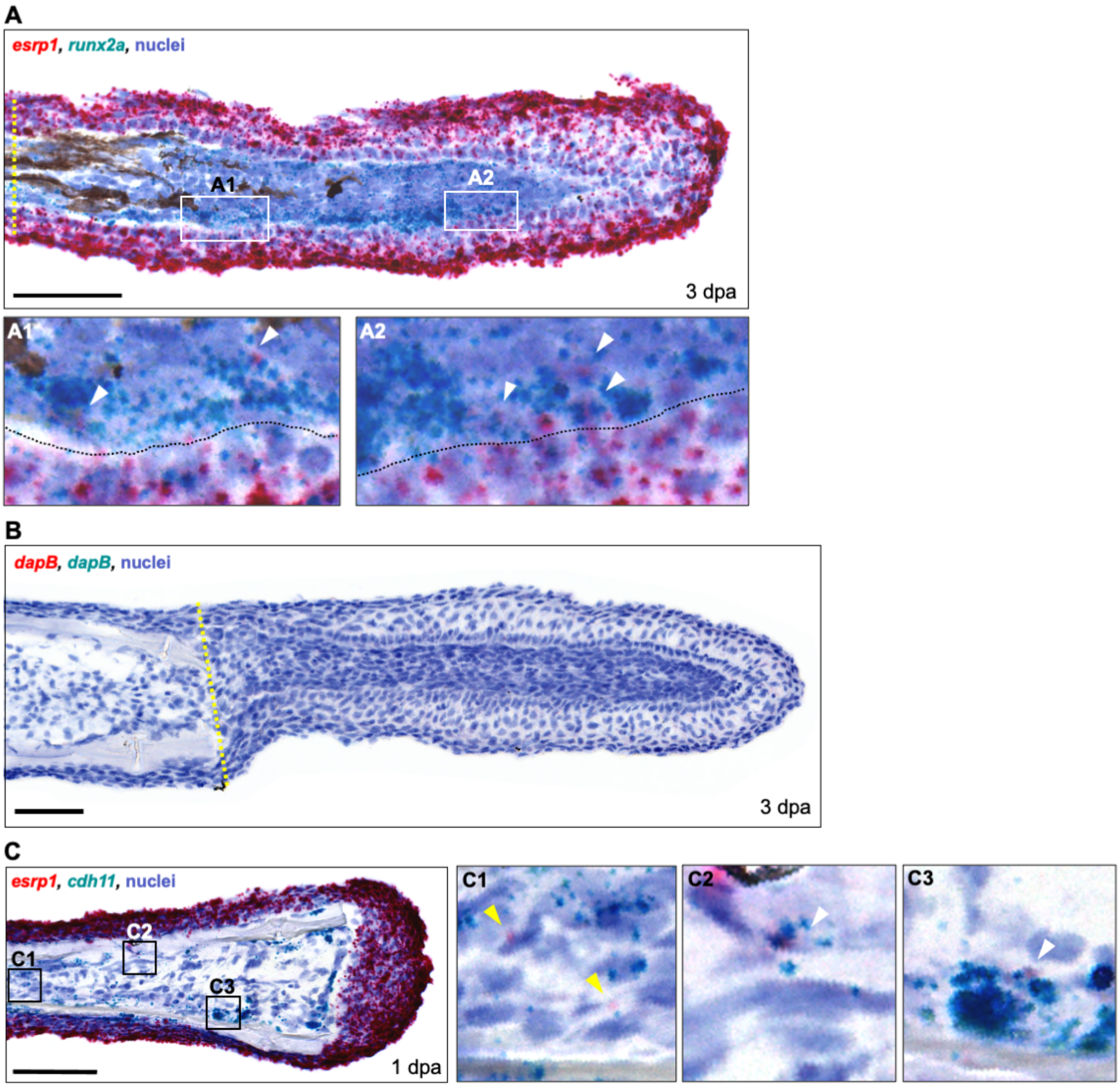


Figure S12. Dual RNA *in situ* hybridization (ISH) in cryosections of regenerating fin tissues at various timepoints, Related to Figure 5. Yellow dashed line indicates the amputation plane. (A) Dual RNA ISH for *esrp1* and *runx2a* in a 3 dpa fin cryosection. Multiple cells (arrowheads in A1, A2) in both proximal and distal locations within the blastema showed costaining for *runx2a* and *esrp1*. Scale bar = 100 μm. (B) Negative control dual RNA ISH in a 3 dpa fin cryosection. Both control probes target *dapB,* a gene found only in bacteria. Scale bar = 100 μm. (C) Expression of *esrp1* and *cdh11* in a 1 dpa fin cryosection. (C1 inset) *esrp1* was infrequently observed in mesenchymal cells, some of which were *cdh11*- (yellow arrowheads). (C2, C3 insets) Several *cdh11*+ cells near the native bone expressed *esrp1* as well (white arrowheads). Scale bar = 60 μm.

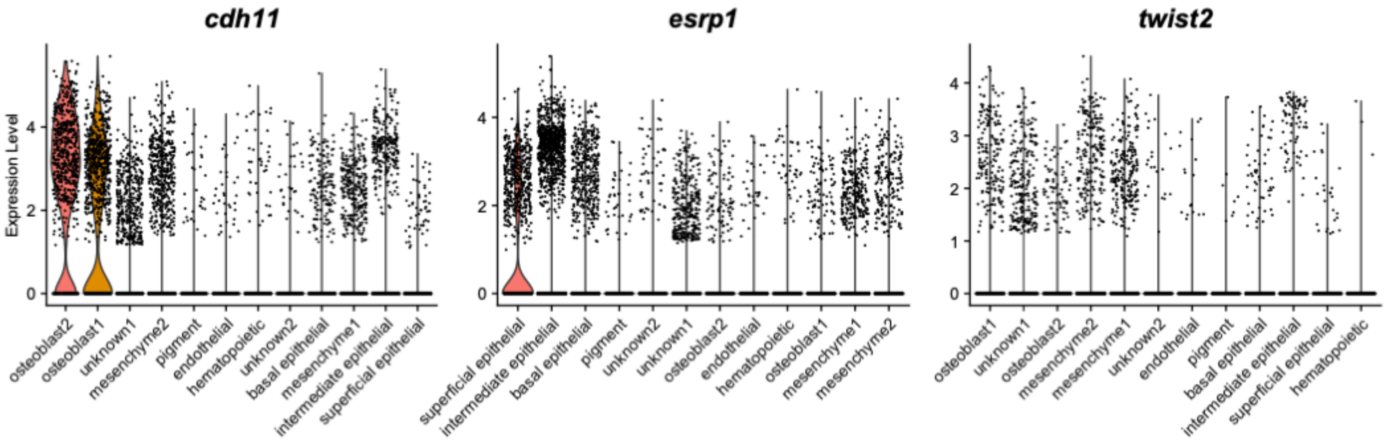


Figure S13. Violin plots visualizing expression levels of *cdh11*, *esrp1*, and *twist2* in the pooled datasets, Related to Figure 5. Clusters are sorted in order of highest to lowest average expression.
